# Dynamic relocalization and divergent expression of a major facilitator carrier subfamily in diatoms

**DOI:** 10.1101/2024.12.16.628690

**Authors:** Shun Liu, Shun-Min Yang, Chris Bowler, Miroslav Obornik, Richard G. Dorrell

## Abstract

Eukaryotic organisms, including microbial members such as protists and green algae, utilise suites of transporter proteins to move essential metabolites across cell organelle membranes. Amongst these different transporter families, the MCF (Mitochondrial Carrier Family) is one of the most diverse, encompassing essential NAD+ and ADP/ ATP translocators, as well as amino acid, sugar and cofactor transporters. They are typically associated with the mitochondrial inner membrane but some display more dynamic localization. Here, we perform a brief census of MCF domains in the genome of the model diatom alga *Phaeodactylum tricornutum*, identifying a new family of three proteins (termed here and elsewhere “MCFc”) with strong internal sequence conservation but limited similarity to other MCF proteins. Considering both phylogenetic data and experimental localization, we posit that MCFc is widespread across algae with complex red chloroplasts alongside some primary green algae, and contains multiple sub-families targeted to diatom mitochondria, plastids, and endomembranes. Finally, using data from *Tara* Oceans, we identify putative roles for MCFc in diatom cells, including a probable association of the plastid-targeted Phatr3_J46742 sub-family in cellular nitrate assimilation. Our data provide insights into the evolutionary diversification of the membrane transport mechanisms associated with diatoms and other eukaryotic algae.

## Introduction

Eukaryotic cells are defined by the presence of discrete organelles (i.e., nuclei, mitochondria, chloroplasts) that direct diverse and essential cellular activities. Many of these organelles date back to the last common eukaryotic ancestor, alongside more recent and lineage-specific organelles acquired through endosymbiosis and through duplication events (Hammond *et al*., 2022; More *et al*., 2024; Novák Vanclová and Dorrell, 2024; Richards *et al*., 2024). As each of these organelles is separated by others by one or multiple hydrophobic membrane bilayers, it is essential that metabolites, proteins and cofactors required for their function, and both metabolic and waste products of their function, are actively transferred into and out of organelles by a diverse range of transporter proteins (Saier, Tran and Barabote, 2006). Protein transporters are highly diverse in terms of their structural evolution, metabolic functions and distribution across the eukaryotic tree of life (Saier, Tran and Barabote, 2006). For example, of nearly 80 membrane transporters associated with the chloroplasts of the model plant *Arabidopsis* only 11 are conserved with the model diatom *Phaeodactylum tricornutum*, and it is likely that even the protein homologues shared between both species have different substrates and kinetic activities despite their underlying structural similarity (Moog *et al*., 2020; Liu *et al*., 2022).

One of the more diverse and enigmatic membrane transporters are the mitochondrial carrier family (MCF) proteins. The MCF proteins (also known as solute carrier family, SLC25; PFAM domain PF00153) are highly diverse, forming the single most expanded protein carrier family in the human genome (48 known genes), and include essential ADP/ ATP transporters, phosphate carriers, and essential transporters for amino acids (aspartate, glutamate), cofactors (thiamine) and sugars (ketoglutarate, citrate) (Kunji, 2004; Ruprecht and Kunji, 2020). MCFs are defined by three successive regions each containing two linked hydrophobic domains, with a consensus sequence each of P-X-[D/E]-X-X-[R/K]_20_ [D/E]GXXXX[W/Y/F][K/R]G (Kuan and Saier, 1993; Kunji, 2004; Monné and Palmieri, 2014). While the majority of annotated MCFs are believed to localize to the mitochondrial inner membrane, some do not, including NAD+ transporters localized to peroxisomes in humans, yeast and plants (Palmieri *et al*., 2001; Agrimi, Russo, Pierri, *et al*., 2012; Toleco *et al*., 2020) and to plant chloroplasts (Palmieri *et al*., 2009) and ATP and other nucleotide triphosphate transporters localized to the chloroplasts of both plants (Ast *et al*., 2009; Nunes-Nesi, Cavalcanti and Fernie, 2020). Beyond different localizations, substantial internal differences exist in the structure and functions of MCF, with many described examples showing less than 10% reciprocal protein sequence similarity to one another (Kuan and Saier, 1993).

Beyond established cell biology models in animals, fungi, and plants, eukaryotes have diverged into a wide range of lifestyle and cell forms (More *et al*., 2024; Penot-Raquin *et al*., 2024). These include diatoms, which are typically unicellular and chain-forming algae abundant both in marine and freshwater habitats, separated from plants by hundreds of millions of years of evolutionary distance, who perform nearly one-quarter of planetary primary production (Serôdio and Lavaud, 2020). Diatom differ from plants in having chloroplasts surrounded by four (not two) membranes, the outermost of which is contiguous with the endoplasmic reticulum (Ast *et al*., 2009; Moog *et al*., 2020), derived from the secondary or more complex endosymbiosis of a eukaryotic red alga itself with a nucleus and chloroplast (Novák Vanclová and Dorrell, 2024). Diatom biology is progressively being understood through cellular models such as *P. tricornutum*, for which extensive synthetic biology techniques and high-throughput gene expression and proteomics data are available (Siaut *et al*., 2007; Liu *et al*., 2022; Huang *et al*., 2024; Villar *et al*., 2024). In addition, reflecting their abundance in marine communities, diatoms are well-represented in environmental sequencing expeditions such as *Tara* Oceans, giving us unprecedented opportunities for identifying the spatial occurrence, abundance and relative expression of the dark matter of their genomes in the wild (Liu *et al*., 2022; Penot-Raquin *et al*., 2024).

Here, concomitant with a mutant study of a novel *P. tricornutum* MCF protein with apparent chloroplast localization (Phatr3_J46742) (Giustini *et al*., 2024) we present an overview of MCF protein diversity, function, and expression trends in diatoms. We note that Phatr3_J46742 forms a novel MCF subfamily including two other members (Phatr3_J42874, Phatr3_J46612) that we localize respectively to the endomembrane system and to the mitochondria. Considering a manually curated phylogeny of these proteins, including inferred localization, from over 250 algal genomes and transcriptomes, we infer this protein likely derives from green algae, with recent paralogy events underpinning the progressive recruitment of Phatr3_J46742 from the mitochondria to the chloroplast (Liu *et al*., 2022). We further consider both cellular (Villar *et al*., 2024) and environmental expression data from diatoms (Penot-Raquin *et al*., 2024) to suggest that these proteins may have diverged in function, alongside changing localization, and play different physiological roles in the diatom cell.

## Materials and Methods

### Phylogenetics

MCF domains (PF00153) were identified in peptide models from the version 3 annotation of the *Phaeodactylum tricornutum* genome using hmmsearch with e-value 10^-05^, per (Kuan and Saier, 1993; Villar *et al*., 2024). *In silico* localizations, KEGG functions, and transcriptional coregulation data were obtained from the DiatomicBase database of *P. tricornutum* genome annotations, per (Liu *et al*., 2022; Villar *et al*., 2024) Fold-enrichments and P-values of enrichment of each protein in plastid-enriched proteome fractions were taken from (Huang *et al*., 2024).

To identify MCFc homologues, the complete sequence of Phatr3_J46742, Phatr3_J42874 and Phatr3_J46612 in the *P. tricornutum* v3 genome annotation (Villar *et al*., 2024) were searched against a composite library of sequences across the tree of life (Liu *et al*., 2022; Penot-Raquin *et al*., 2024). This library consisted of a complete version of uniprot (accessed 2018), along with decontaminated versions of the MMETSP transcriptome project, and JGI PhycoCosm algal genomes, as previously described (Keeling *et al*., 2014; Suzek *et al*., 2015; Grigoriev *et al*., 2021). Possible homologues were extracted by reciprocal BLASTp best hit, with threshold e-value for the initial BLAST search 1×10^-05^, and best hits identified by reciprocal search against the entire *P. tricornutum* genome (Liu *et al*., 2022).

Probable localisations of sequences derived from lineages with four membrane plastid structures (cryptomonads, haptophytes and ochrophytes) were inferred using the consensus predictions of SignalP v 3.0, ASAFind, HECTAR, and MitoFates (Bendtsen *et al*., 2004; Gschloessl, Guermeur and Cock, 2008; Fukasawa *et al*., 2015; Gruber *et al*., 2015); as previously described (Liu *et al*., 2022). A taxonomically representative subset of the sequences obtained were aligned using MAFFT v 5.0, MUSCLE v 8.0, and GeneIOUS v 4.76 (Edgar, 2004; Kearse *et al*., 2012; Katoh, Rozewicki and Yamada, 2019); trimmed manually and with trimAl in –gt 0.5 setting (Capella-Gutiérrez, Silla-Martínez and Gabaldón, 2009); and consensus phylogenies were inferred using MrBayes and RAxML programmes integrated into the CIPRES server, using three substitution matrices (GTR, Jones/ JTT, and WAG), as previously described (Huelsenbeck and Ronquist, 2001; Miller, Pfeiffer and Schwartz, 2010; Stamatakis, 2014).

### Experimental localization

The complete sequences of Phatr3_J42874 and Phatr3_J46612, as amplified from cDNA and using *P. tricornutum* version 3 genome annotation gene models, were cloned into a pPhat-Fcp::eGFP-SHBLE vector by Gibson cloning, using previously defined methodology(Ast *et al*., 2009; Gile *et al*., 2015), yielding constitutively overexpressed constructs with a C-terminal GFP tag. For Phatr3_J46742, the complete coding sequence was found to be unable to produce GFP fluorescence, so a truncated construct, consisting of the last 209 bp of the predicted 5’ UTR and first encoded 261 bp (87 aa) of the coding sequence was similarly cloned. This construct was designed both to cover several potential alternative initiator codons (in-frame methionines) in the CDS 5’ end, but to avoid out-of-frame initiator codons that could interfere with the correct translation of the C-terminal GFP fusion. Vector sequences confirmed by PCR and Sanger sequencing were introduced into wild-type *P. tricornutum* strain Pt1 by biolistic transformation, following the methodology of (Falciatore *et al*., 1999).

Transformant colonies were selected first on f/2-agar plates supplemented with 100µg ml^-1^ zeocin. Resistant colonies were transferred to liquid media, and GFP expression was first assessed by Western blot using a mouse anti-GFP antibody (ThermoFisher) (Ast *et al*., 2009). Constructs founds to express GFP were visualised by confocal microscopy. Mitochondria in cell lines were stained with MitoTracker Orange (Sigma) (1µg ml^-1^) for 30 minutes prior to visualization, following (Tanaka *et al*., 2015). Typically, fluorescence was assessed using a Leica SP8 confocal microscopy (IBENS imaging platform), with a 485 nm excitation laser and detection window 500-525 nm to identify GFP fluorescence, and 548 nm excitation laser and detection window 565-590 nm to identify MitoTracker fluorescence, chlorophyll autofluorescence detected over a 650-700 nm window and whole cells visualized with bright-field microscopy. Negative controls (either MitoTracker-stained wild-type cells, or unstained GFP expressing cells) were used to confirm specificity of each detection window and set exposure thresholds, and fluorescence was confirmed via sequential excitation of each laser to exclude the possibility of crosstalk between GFP and Mitotracker fluorescence (**Fig. S7**). Composite fluorescence images were prepared with ImageJ.

### Gene expression analysis

Gene coregulation analysis in *P. tricornutum* was performed using methodology previously described in (Liu *et al*., 2022; Penot-Raquin *et al*., 2024), using two gene expression databases (DiatomPortal-microarray; PhaeoNet-RNAseq) converted into ranked values to allow calculation of Spearman correlation coefficients. Core organelle metabolic pathways for the plastid and mitochondria were constructed from previously available models (Kroth *et al*., 2008; Ast *et al*., 2009; Smith *et al*., 2019).

*Tara* Oceans calculations were performed following previously defined methodology (Liu *et al*., 2022) based on phylogenetic reconciliation of environmental sequences to annotated cultured species homologues. Briefly, potential meta-transcriptome (“metaT”) and meta-genome (“metaG”) homologues of MCFc proteins were identified by BLASTp with threshold e-value 10^-05^ in *Tara* Oceans meta-gene data (Busseni *et al*., 2019; Penot-Raquin *et al*., 2024). These were searched by BLASTp against the complete version 3 *P. tricornutum* genome and against the previously generated cultured species alignment, with only sequences that retrieved *(i)* an MCFc family gene (Phatr3_J42874, Phatr3_J46612, Phatr3_J46742) and *(ii)* a cultured diatom as best hit retained for further analysis. Identified putative homologues were aligned against the cultured species MCFc alignment using MAFFT as above, and inspected using RAxML using 350 bootstrap replicates and the PROTGAMMAJTT substitution model. Sequences that reconciled as monophyletic group with each diatom MCFc subfamily, to the exclusion of other proteins and organisms, were assigned as such. Relative metaT and metaG abundances expressed as a fraction of all meta-gene coverage at each station, in both 5-20 and 20-180 µm size fractions at surface and deep chlorophyll maximum (“DCM”) depths; were compared to one another and to different environmental variables via a matrix determinant test, following (Liu *et al*., 2022).

## Results and Discussion

### Diversity of MCF proteins in the Phaeodactylum tricornutum genome includes a novel sub-family

First, we constructed a census of MCF domains encoded in the version 3 annotation of the *P. tricornutum* genome, identified using hmmsearch of the associated PFAM (PF00153) using hmmsearch with e-value 10^-05^ (**Fig. 1**) (Liu *et al*., 2022; Villar *et al*., 2024). A total of 65 genes encoding potential MCF domains were identified, slightly more than those found in the genomes of *Arabidopsis thaliana* (59) or the model green alga *Chlamydomonas reinhardtii* (46) (Fernie, Cavalcanti and Nunes-Nesi, 2020). We note that amongst published diatom genomes (e.g., housed in JGI Phycocosm) and transcriptomes (MMETSP) the number of annotated MCFs ranges from 22 (*Chaetoceros brevis* CCMP164 MMETSP transcriptome) to 113 (*Corethron pennatum* MMETSP transcriptome), which broadly corresponds to the range (39-141) associated with sequenced plant genomes (**Fig. S1**) (Keeling *et al*., 2014; Grigoriev *et al*., 2021). Through gene coregulation module, *in silico* localization prediction, PFAM and Kofam annotations (Villar *et al*., 2024) and also pairwise reciprocal BLASTp similarities, we attempt a preliminary classification of *P. tricornutum* MCF proteins based purely on primary sequence in **Fig. 1**. A searchable version of this figure is provided as **Table S1**, sheet 1.

**Fig. 1.**
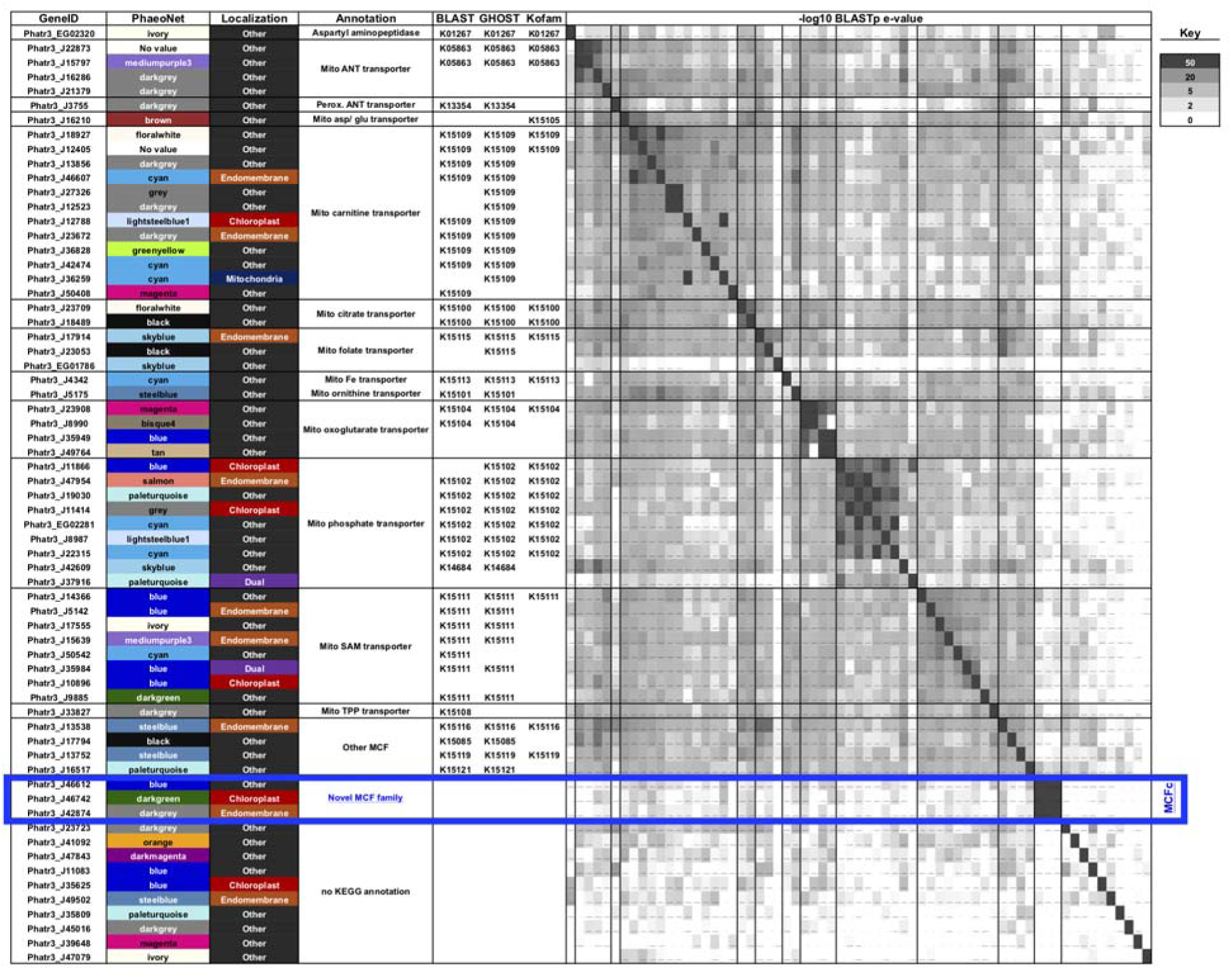
Diversity of MCF domain proteins in *Phaeodactylum tricornutum.* This figure shows all 65 genes in the *P. tricornutum* genome version 3 annotation inferred to encode a Mitochondrial Carrier Family (PF00153.30) PFAM (Palmieri *et al*., 2011), alongside a heatmap of reciprocal BLASTp x BLASTp –log10 evalue. WGCNA Transcriptional modules (from (Ait-Mohamed *et al*., 2020)), consensus localization predictions, and KEGG annotations are provided per (Villar *et al*., 2024). Genes are grouped by inferred function, with a novel MCF sub-family (MCFc: Phatr3_J42874, Phatr3_J46612, Phatr3_J46742) with high (e< 10-50) reciprocal BLASTp similarity to one another, but no described KEGG function, highlighted by a blue box.

Considering KEGG annotations, the *P. tricornutum* genome encodes multiple probable adenine nucleotide transporters with homology both to human mitochondrial (KEGG ID: K05863; Phatr3_J22873, Phatr3_J15797, PhatrJ_16286, Phatr3_J21789) and peroxisomal proteins (KEGG ID: K13354, Phatr3_J3755) (Neckelmann *et al*., 1987; Agrimi, Russo, Scarcia, *et al*., 2012), although we note that the *in silico* targeting predictions of the *P. tricornutum* homologues were unclear. Further MCF transporters were annotated by KEGG to be homologues of known aspartate/ glutamate, carnitine/ acylcarnitine, folate, iron, ornithine, oxoglutarate, phosphate, S-adenosylmethionine and thiamine pyrophosphate transporters (Palmieri, 2004) (**Fig. 1**). Several of these gene families were highly expanded (e.g., twelve protein homologues to carnitine/ acylcarnitine transporters) and in many cases showed strong homolgy (reciprocal BLASTp evalue < 10^-20^) to one another (**Fig. 1**). Surprisingly, only two (Phatr3_J36259, a putative mitochondrial carnitine transporter; and Phatr3_J49674, an oxoglutarate transporter) had N-terminal mitochondrial presequences detectable by targeting predictors (HECTAR, which incorporates MitoProt2 iPSort, Predotar, PredSL, and TargetP; and MitoFates) typically able to detect diatom mitoproteins with high sensitivity (Gile *et al*., 2015; Hammond *et al*., 2022). Two putative phosphate transporters, Phatr3_J11414 and Phatr3_J37916, and one S-adenosylmethionine transporter, Phatr3_J35894 were inferred to possess dual-targeting sequences (i.e., presequences hydrophobic enough to be recognised by SignalP containing a post-cleavage ASAFAP motif; but hydrophilic enough to also function as a cleavable transit peptide) that might direct them both to the plastid and mitochondria (**Fig. 1**). While dual-targeting is known in diatom organelles it is typically associated with proteins involved in translation (e.g., aminoacyl-tRNA synthetases) and the degree to which it is associated with transporter proteins awaits experimental study (Gile *et al*., 2015; Liu *et al*., 2022). It remains to be determined if any of the remaining 55 MCF proteins that lack identifiable mitochondrial presequences are targeted to *P. tricornutum* mitochondria by alternative means, e.g. internal C-terminal targeting sequences (Diekert *et al*., 1999; Lee *et al*., 1999), particularly given they are typically associated with the inner mitochondrial membrane which is viewed largely not to be accessible to direct peptide insertion (Palmieri *et al*., 2011; Hammond *et al*., 2022). An alternative possibility may be the presence of cryptic exons encoding N-terminal targeting sequences in the annotated 5’ UTRs of these genes, which will be best verified by experimental confirmation of each gene model (e.g., using 5’ RACE) (Russo *et al*., 2015)

More surprisingly, we detected 14 MCF proteins with identifiable signal peptides (e.g., using SignalP 5.0), suggesting endomembrane localization. Five of these, including Phatr3_J46742, contained bipartite signal and transit peptides (inferred either using ASAFind or HECTAR prediction) associated with plastid-targeting presequences in diatoms (Moog *et al*., 2020; Liu *et al*., 2022). Amongst these putatively plastid-targeted MCFs were an inferred carnitine transporter (Phatr3_J12788), a phosphate transporter (Phatr3_J11866) and an S-adenosylmethionine transporter (Phatr3_J10896) detected in previous analyses of the *P. tricornutum* plastid transporter proteome (Liu *et al*., 2022). Of note, all three of these transporters appear to be co-expressed in published *P. tricornutum* gene expression data with chloroplast light harvesting complex proteins (Phatr3_J12788) and photosynthesis and Calvin cycle metabolism (Phatr3_J10896; Phatr3_J11866), supporting likely plastidial functions (Villar *et al*., 2024). Considering a previously published experimental proteomic dataset from *P. tricornutum* (Huang *et al*., 2024), all of the four putatively plastid-targeted MCF transporters found at detectable levels (Phatr3_J10896, Phatr3_J11886, Phatr3_J35625, Phatr3_J46742), were enriched in plastid over whole-cell fractions, coherent with the *in silico* targeting predictions (**Fig. S1**). That said, the majority of the MCF domain proteins, irrespective of localization prediction, were enriched in plastidial fractions, with two putatively endomembrane-targeted transporters (Phatr3_J13538, Phatr3_J47954) and four that lack any recognizable targeting presequences (Phatr3_J19030, Phatr3_J23709, Phatr3_J23908, Phatr3_J35949) deemed to be significantly enriched (P < 0.01; **Fig. S1**) (Huang *et al*., 2024). The occurrence of these presumably non-plastidial transporters in the plastid-enriched fraction may relate to the close association of the *Phaeodactylum* plastids, endomembrane, and indeed mitochondria in vivo, which may impede separation of their associated proteomes by organelle enrichment approaches (Schober *et al*., 2019; Uwizeye *et al*., 2021).

Finally, alongside Phatr3_J46742, we note that two additional MCF transporters (Phatr3_J42874, Phatr3_J46612) showed high reciprocal BLASTp similarity (e-values of < 10^-60^ in each pairwise combination) but limited similarity (P > 10^-05^) to any other described MCF in the *P. tricornutum* genome (**Fig. 1**). These MCFs further lacked any apparent KEGG annotations, considering combined BLASTkoala, GHOSTkoala, and KofamKoala annotations (Jin *et al*., 2023; Kanehisa *et al*., 2024) and may represent a new functional subfamily of MCFs. Phatr3_J42874 was predicted to encode a signal peptide but lacked a downstream ASAFAP site or transit peptide, consistent with endomembrane localization, and Phatr3_J46612 lacked any apparent targeting presequences. Consistent with nomenclature used in a simultaneous mutant study of Phatr3_J46742 (Giustini *et al*., 2024), we herefore refer to this new MCF family as “MCFc”, i.e. “mitochondrial carrier family chloroplastic”.

### Phylogeny of MCFc reveals homologues in other complex red plastid-bearing and primary green algae

To reconstruct the deeper evolutionary history of MCFc, we searched for homologues of each complete *P. tricornutum* peptide sequence across a composite genome and transcriptome library of the entire tree of life, following previously defined methodology (Liu *et al*., 2022). Briefly, this involved performing a primary BLASTp of each MCFc subunits, with threshold evalue of 10^-05^ against a composite library of over 250 diatom and related algal genomes and transcriptomes, alongside complete copies of uniref (Suzek *et al*., 2015), MMETSP transcriptomes (Keeling *et al*., 2014) and JGI Phycocosm genomes (Grigoriev *et al*., 2021), followed by a reciprocal BLASTp search against the version 3 annotation of the *P. tricornutum* genome (Villar *et al*., 2024). Only sequences that yielded an MCFc family protein as a BLAST best hit were retained for downstream analysis. From this approach, we identified nearly 2000 potential homologous sequences (**Fig. S2**; **Table S1**). These homologues were distributed across a wide range of eukaryotic groups, although were primarily found in lineages with a history of complex endosymbiosis (i.e., cryptomonads, haptophytes, ochrophytes and dinoflagellates) (Novák Vanclová and Dorrell, 2024; Penot-Raquin *et al*., 2024); and were noticeably absent from prokaryotes (**Fig. S2A**). Considering probable localisations, the vast majority of chloroplast-targeted sequences in species with four membrane-bound plastids were found to either be homologous to Phatr3_J46742 (in diatoms and cryptomonads), or to Phatr3_J42874 (in other ochrophytes and in haptophytes), with Phatr3_ J46612 homologues largely lacking clear localisations (**Fig. S2B**).

Next, we constructed a 78 taxa x 262 aa phylogeny of a representative subset of the homologues identified, which was manually trimmed to remove poorly aligned sequences and uninformative regions such as N-terminal presequences (**Table S1**, sheets 2-6). This phylogeny revealed a complex evolutionary history of MCFc family proteins (**Fig. 2**). Phatr3_J46742 formed a robustly supported (MrBayes PP = 1.0; RAxML bootstrap support > 80%; 3 substitution matrices), monophyletic group with other diatom chloroplast-targeted proteins, consistent with its apparent proteomic localisation. This formed a sister-clade to the predominantly endomembrane-targeted proteins related to Phatr3_J42874, and a large clade of dinoflagellate, cryptomonad and haptophyte sequences, which might be consistent with endosymbiotic or other transfers of genes encoding plastid-targeted proteins from ochrophytes into these lineages (Liu *et al*., 2022; Novák Vanclová and Dorrell, 2024). Notably, the closest sister-group of the combined clade of ochrophyte, cryptomonad, haptophyte and dinoflagellate sequences was a monophyletic clade of prasinophyte green algae, including members of the Mamiellophyceae, Pyramimonadales, Prasinococcales and Chlorodendrales (**Fig. 2**). This may indicate that the MCFc gene family is of green origin, although this hypothesis requires further testing with more deeply-positioned sister-groups.

**Fig. 2.**
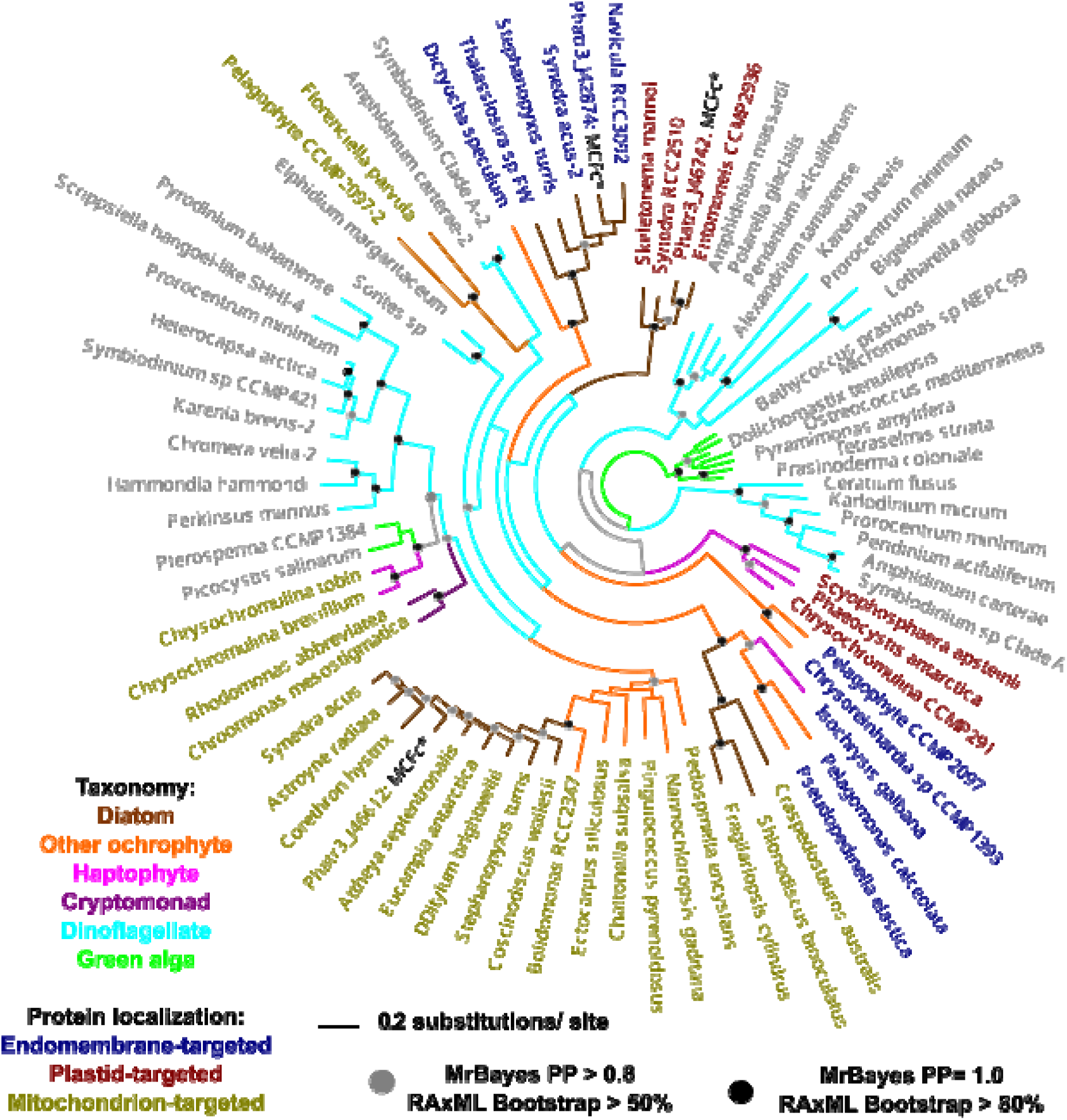
Phylogeny of subselected MCFc proteins across the tree of life. The topology shown is the consensus of MrBayes consensus trees and RAxML best-scoring trees, each realised with three substitution matrices (GTR, Jones/ JTT, WAG). Branches are shaded by taxonomy, and leaf nodes by predicted in silico localization based on consensus targeting predictions (HECTAR, ASAFind, SignalP, MitoFates) and experimental localization with GFP (*P. tricornutum* proteins only). *P. tricornutum* proteins are highlighted with the appellation “MCFc*”. Nodes are labelled with coloured circles that define minimal MrBayes posterior probabilities and RAxML bootstrap values across all substitution matrices studied. Broadly, three clades of proteins are visible in algae with chloroplasts of secondary or higher red origin, each exemplified by *P. tricornutum* proteins, with respectively plastid (Phatr3_J46742), endomembrane (Phatr3_J42874) or mitochondrial (Phatr3_J46612) localization, with a green algal sister-group.

### MCFc homologues show different subcellular localizations suggestive of evolutionary history

Next, we localized all three MCFc family proteins in *P. tricornutum* cells using C-terminal GFP fusion constructsand confocal microscopy (**Fig. 3**, **Table S2**). Phatr3_J42874 and Phatr3_J46612 were localized using full-length constructs, while Phatr3_J46742 was localized using a short N-terminal region consisting of the final 209 bp of the 5’ UTR and the first encoded 87 aa, which was found to produce strong GFP expression. The Phatr3_J46742 N-terminal region clearly localizes within the chloroplast stroma, and does not colocalize with the mitochondria, as assessed by staining with MitoTracker Orange (**Fig. 3**, panel A). While we cannot exclude that the specific localization within the stroma is artifactual (e.g., due to using an N-terminal rather than full-length construct driven by a constitutive and strong promoter (Gile *et al*., 2015; Moog *et al*., 2020)) it is notably supported by experimental proteomic data above, where Phatr3_46742 shows a strong enrichment (14.7-fold, P = 0.12) in plastid-enriched to whole-cell fractions **(Fig. S1**) (Huang *et al*., 2024). The remaining two MCFc transporters did not localize to plastids. Phatr3_J42874 was found in contrast to have a filamentous extra-plastidial localization that did not overlap with the mitochondria, which may suggest endomembrane targeting in agreement with *in silico* localization predictions. In contrast, Phatr3_J46612 clearly overlapped with the MitoTracker signal, confirming it as a bona fide mitochondrial transporter (**Fig. 3**, panels B to C). Overall, our data are consistent with the MCFc family having both plastidial and non-plastidial members. Considering the phylogenetic proximity of Phatr3_J46742 to Phatr3_J42874, and comparative distance to Phatr3_J46612, we propose it is likely to have been an ancestrally mitochondria-targeted protein (similar to Phatr3_J46612) that was subsequently recruited to the plastid, potentially via an endomembrane-targeted intermediate (Phatr3_J42874) (**Fig. 2**).

**Fig. 3.**
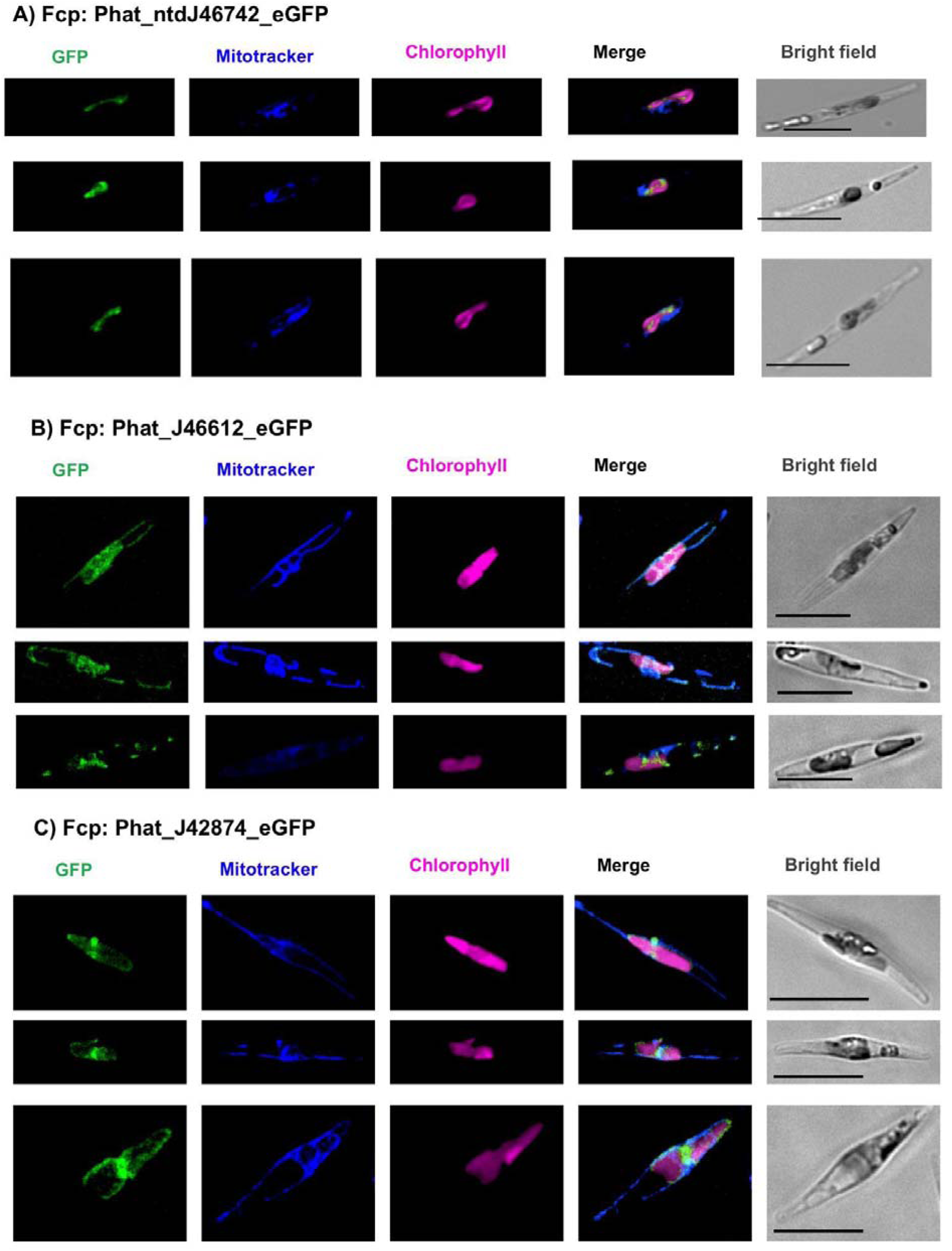
Differential localization of *P. tricornutum* MCFc homologues. This figure shows confocal microscopy images of transgenic *P. tricornutum* expressing C-terminal linked GFP constructs for three closely-related MCFc transporters, Phatr3_J46742 (A), Phatr3_J46612 (B), and Phatr3_J42874 (C). Phatr3_J46612 and Phatr3_J42874 are full-length constructs, and Phatr3_J46742 is an N-terminal construct avoiding out-of-frame internal ORFs that may otherwise impede in-frame GFP translation. All cell lines are stained with MitoTracker Orange (false-stained blue) per Tanaka et al., 2015. Control images (stained and unstained cell lines expressing unlinked cytoplasmic GFP, and Mitotracker-stained wild-type cell lines) are shown in **Fig. S6**. Scale bar for all images = 10 μm.

### Gene expression analysis of MCFc genes in P. tricornutum and Tara Oceans data suggest discrete biological functions

Given the distinct localizations of different MCFc family proteins in *P. tricornutum*, we considered whether they are likely to perform linked or separate transport functions. First, we used a previously published approach in which we performed Spearman correlations of ranked transcript relative abundance between each MCF family protein gene (Phatr3_J46742, Phatr3_J42874, Phatr3_J46612) across a large dataset of over 300 normalized and previously published microarray and RNAseq libraries (Ashworth *et al*., 2016; Liu *et al*., 2022; Penot-Raquin *et al*., 2024; Villar *et al*., 2024). The data from these analyses are provided as a pivot table (**Table S3**).

Briefly, the plastid localized MCFc Phatr3_J46742 showed typically weak or negative correlation to most genes encoding core *P. tricornutum* organelle metabolic processes, including both other MCFc transporters (Spearman correlation < −0.2; **Fig. S3A**; **Table S3**, sheets 1-3). This may indicate that this gene does not form a core component of organelle metabolism, and may be inducibly expressed under certain conditions only. Phatr3_J46742 nonetheless shows strong positive coregulation (Spearman correlation > 0.5; **Fig. S3B**) with a small number (17) of genes encoding mitochondria- and/ or plastid-targeted proteins with identifiable (KEGG, PFAM functions). These notably include a mitochondria-targeted major facilitator superfamily transporter (geneID : Phatr3_J27361) which is strongly positively coexpressed (correlation 0.706). Across a broader search of core organelle metabolic pathways (**Table S3,** sheets 4-5) we also identified strong positive coregulation within the plastid (correlation > 0.45) to both phosphofructokinase 1 (PFK) and fructose-1,6-bisphosphatase (FBP) involved in the Calvin cycle, and a strong negative coregulation (correlation <-0.5) to glutamate synthase (GOGAT) implicated in nitrogen fixation, which may point to further interacting partners in plastid metabolism (Kroth *et al*., 2008; Smith *et al*., 2019).Phatr3_J42874 and Phatr3_J46612 showed in contrast strong positive coregulation with a wide range of core plastid- and mitochondria-targeted proteins, with in particular (one-way ANOVA, P < 0.01) larger positive coregulation values observed for mitochondria-targeted glycolytic and TCA cycle proteins (Phatr3_J42874) and plastidial ribosome and carbon concentrating mechanism proteins, and mitochondrial complex I (Phatr3_J46612) than other proteins targeted to the same organelle (**Table S3,** sheets 6-9).

Next, we considered under what discrete conditions different MCFc family genes were differentially expressed in *P. tricornutum*, leveraging normalized RNAseq data integrated into the DiatomicBase server (Villar *et al*., 2024). We noted that each MCFc gene showed responses to different environmental stimuli, potentially suggesting different physiological roles (**Fig. S4**). Phatr3_J46742 was found to be upregulated in nitrate reductase knockout lines compared to control lines, as well as in knockout lines for the blue light receptor aureochrome after 60 minutes illumination with red or blue light, suggestive of roles both in nitrate metabolism and photoperception (McCarthy *et al*., 2017; Mann *et al*., 2020). It was conversely downregulated both under CO_2_ and phosphate depletion conditions, which might either relate to linked functions to these nutrients or changes in nutrient stoichiometry (e.g., N to P ratio) in these cells (Li *et al*., 2022; Shu *et al*., 2022). In contrast, Phatr3_J42874 was found to be upregulated under phosphate depletion, and in aureochrome knockout lines only under very short-term (10 minute) illumination conditions; while Phatr3_J46612 was downregulated both in aureochrome knockouts, and knockdown lines for the mitochondrial alternative oxidase (Murik *et al*., 2019) (**Fig. S4**).

Finally, we searched meta-gene data from *Tara* Oceans for genes encoding each MCF transporter subfamily. This analysis, adapted from (Liu *et al*., 2022), involved retrieving all *Tara* Oceans meta-genes with known MCF annotations, and then identifying those that corresponded to each diatom MCF sub-family by reciprocal BLASTp against the *P. tricornutum* genome and against the curated MCF homologue alignment used for the single-gene tree. The meta-genes that were identified as encoding MCFc sequences from the reciprocal BLASTp search to *P. tricornutum*, and of diatom origin by BLASTp search against the cultured species alignment, were added to the cultured species alignment and used to build a single-gene RAxML tree. Finally, meta-genes that resolved in monophyletic clades with each *P. tricornutum* MCFc protein (Phatr3_J46742, Phatr3_J46612, or Phatr3_J42874) to the exclusion of all non-diatom proteins were annotated as such. Identified meta-genes, their relative abundances in meta-transcriptome (metaT) and meta-genome (metaG) data, and their correlations to measured *Tara* environmental variables are shown in **Table S4**.

Using these data, we could see ubiquitous expression of Phatr3_J46742 meta-genes in metaT data, with particular concentrations in Southern Ocean, Southern Atlantic and coastal mid-Pacific (e.g., Marquesas Islands) sites (**Fig. 4A**). While the same sequences were also abundant in Southern Ocean stations in metaG data, suggesting underlying high abundance of the diatoms encoding these genes, the remaining stations in which we observed high metaT abundances did not have corresponding high metaG abundances (**Fig. 4B**), suggesting transcriptional regulation of Phatr3_J46742 genes in the wild. Taking into account that the stations in which Phatr3_J46742 shows high metaT abundances are typically from Fe-limited but N-replete regions of the global ocean (Busseni *et al*., 2019; Ustick *et al*., 2021), we considered the possibility that nitrogen availability informs Phatr3_J46742 expression levels in *Tara* Oceans data. Strikingly, Phatr3_J46742 metaT abundances show strong positive correlations (*r*^2^ 0.34-0.90, depending on depth and size fraction) to nitrate (NO_3_^-^) concentrations, both considering direct measurements and simulated concentrations based on the Darwin model at 5m depth (**Figs. 5, S5**) (Pesant *et al*., 2015). In contrast, only weak correlations were found between nitrate concentration and metaG relative abundance, with *r*^2^ < 0.2 in all depth and size fractions except the 5-20 µm surface fraction (**Fig. S5**). Moreover, Phatr3_J46742 metaT abundances showed nil (*r*^2^ < 0.05) or even moderate negative correlations to simulated ammonium concentrations in *Tara* Oceans data, which suggests specific nitrate regulation of its expression (**Fig. S6**). A global search of all *Tara* Oceans environmental variables, comparing the strength of correlation to Phatr3_J46742 metaT versus metaG relative abundance via matrix determinant testing, further revealed significant positive relationships between phosphate and silicate abundance on Phatr3_J46742 relative metaT abundance, and negative relationships to PIC (particulate inorganic carbon), suggesting other environmental factors may determine its expression (**Fig. 5**).

**Fig. 4.**
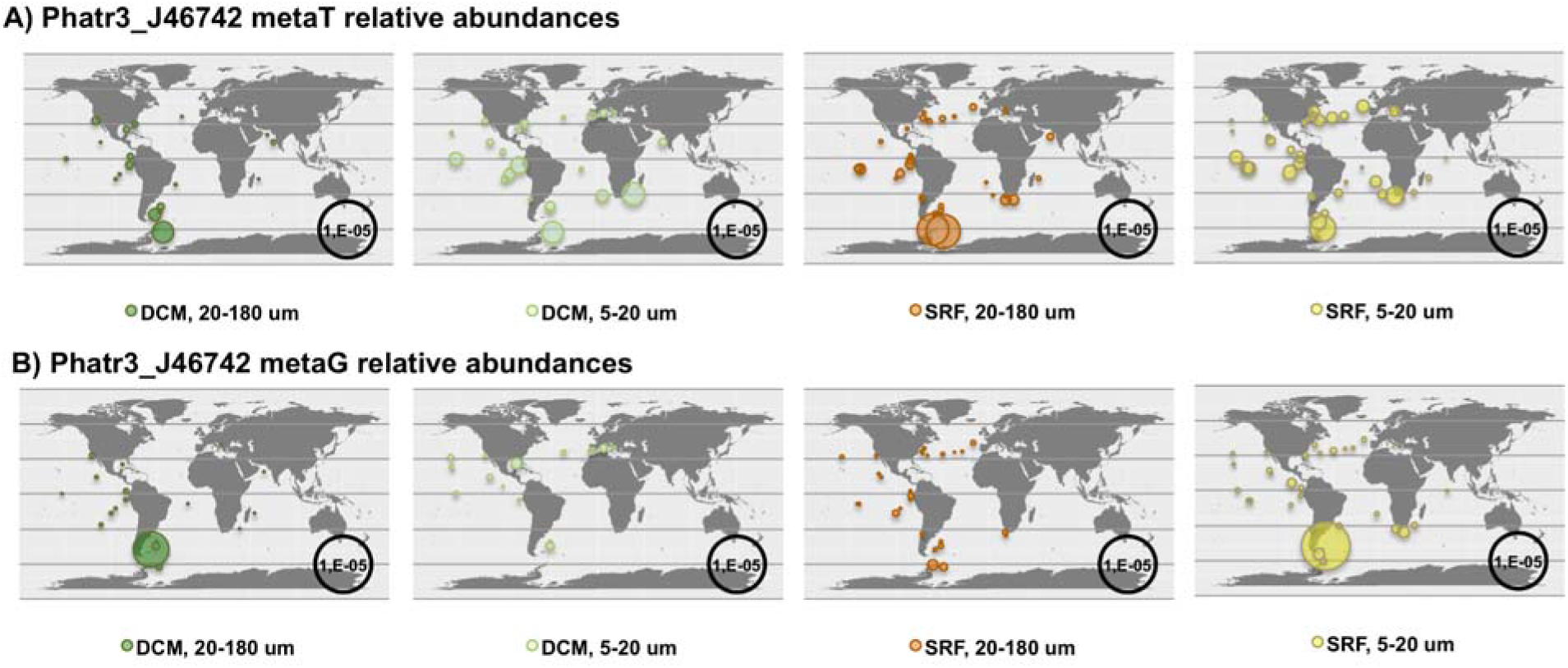
Tara Oceans maps of Phatr3_J46742 relative distributions. This figure shows maps of the relative abundance of MATOUs phylogenetically reconciled to belong to the Phatr3_J46742/ MCFc subfamily of MCF proteins and of diatom origin, based on metaT (meta-transcriptome, **A**) and metaG (meta-genome, **B**) relative mapped abundances, for two depths (DCM-Deep Chlorophyll Maximum; SRF-surface) and size fractions (5-20 μm, 20-180 μm) in which diatoms predominate. Phatr3_J46742 meta-transcripts show cosmopolitan distributions but with particular occurrence in Southern Ocean sites in which their corresponding meta-genes are also abundant, alongside coastal South Atlantic and Pacific sites in which they are not.

**Fig. 5.**
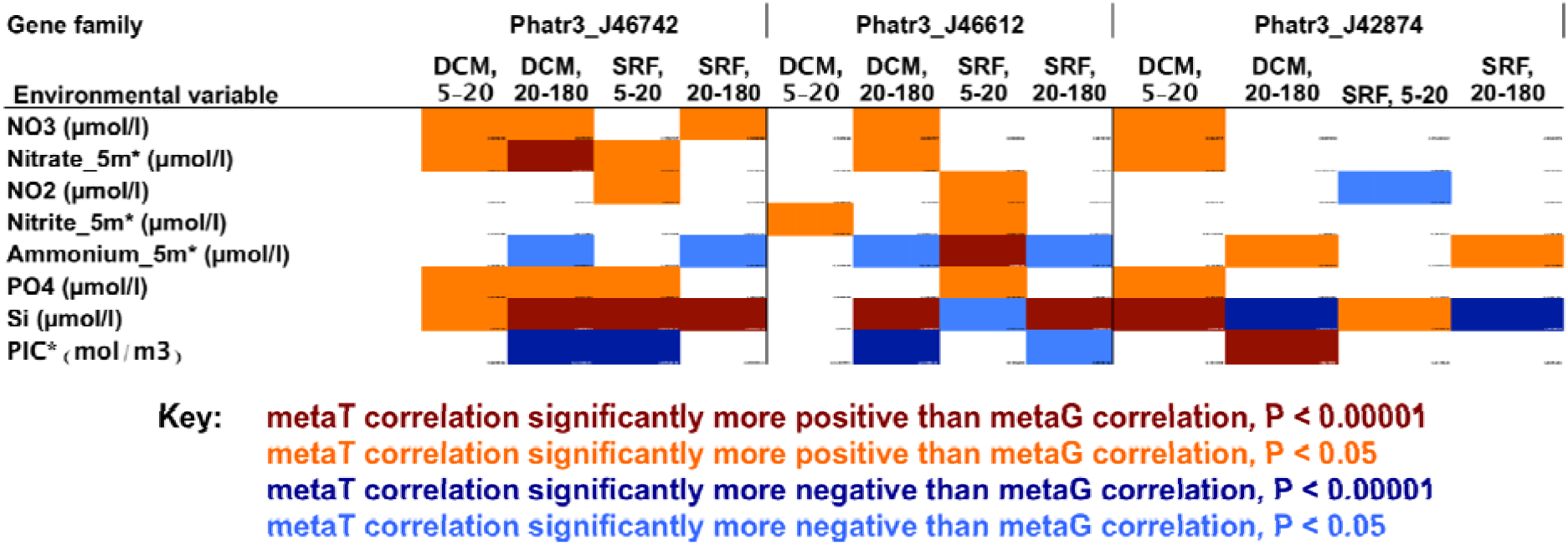
Heatmap of P-values of parametric correlation tests performed between the metaT and metaG total relative abundances of three closely related diatom MCFc transporte families (Phatr3_J46742-chloroplastic; Phatr3_J46612-mitochondrial, Phatr3_J42874-endomembrane) for 5-20 μm and 20-180 μm size fractions and surface and DCM depths. Briefly, these plots show if for each depth and size combination if meta-transcriptome relative abundances show significantly more positive (stronger positive, or weaker negative) correlation to the variable than meta-genome abundances do, accounting for the dependent relationships between metaT and metaG data. Complete data are shown in **Table S4**, sheet 12. Phatr3_46742 shows significantly stronger positive correlations to measured nitrate, phosphate and silicate concentrations in metaT than metaG data, no clear correlations to nitrite, and significantly more negative concentrations to ammonium and to PIC (particulate organic carbon). The same trends are observed much more weakly, or not at all, for the other two MCFc subfamilies.

Meta-genes corresponding to both Phatr3_J42874 and Phatr3_J46612 were both more abundant (in metaG) and more highly expressed (in metaT) in nutrient-rich but iron-limited stations from the Southern Ocean, as typical for diatom transporter proteins (Liu *et al*., 2022). Consistent with roles in primary production and carbon fixation, both Phatr3_J42874 and Phatr3_J46612 showed positive associations between metaT relative abundance and pigment (chlorophyll c3, fucoxanthin) and carbon (carbon total, HCO3-) concentrations, but only in the 5-20 µm size fraction from surface samples (**Table S4**, sheet 12). Of note, for the 20-180 µm size fraction, Phatr3_J46612 like Phatr3_J46742 showed a significantly lower (less positive, or more negative) correlation of metaT abundance compared to metaG abundance against ammonium concentration, whereas the converse (i.e., a significantly more positive, or less negative difference between metaT and metaG correlations) was observed for Phatr3_J42874, which may suggest again separate biological functions (**Fig. 5; Table S4**, sheet 12).

## Concluding Remarks

In this study, we have used evolutionary (BLASTp, phylogeny), experimental (sub-cellular localization), and gene expression approaches (in *Phaeodactylum tricornutum* and *Tara* Oceans) to better understand a novel family of MCF proteins, termed MCFc, in diatom algae. These data broadly suggest that MCFc stands apart from other MCF protein families with known functions,is predominantly restricted to primary green algae and algae with complex red chloroplasts, and contains distinct sub-families with probable mitochondrial, endomembrane, and plastid localizations. These sub-families do not however have similar transcriptional patterns to one another, showing co-regulation to different components of *P. tricornutum* core mitochondrial and plastid metabolism, and showing different expression responses to nitrogen and carbon in *Tara* Oceans. The consistent response of Phatr3_J46742, indicative of the chloroplast-targeted sub-family to nitrate concentrations in metaT but not metaG data, despite nil or negative correlations to ammonium concentrations, may suggest associative roles in nitrate uptake, reduction and assimilation (Busseni *et al*., 2019; Smith *et al*., 2019). These roles, and indeed the functions of other MCFc family transporters, will be best explored via the interrogation of complemented knockout mutants in model species such as *P. tricornutum*, as explored in an independent study of this gene (Giustini *et al*., 2024), alongside *in vitro* identification of substrates and kinetic profiling for each transporter (Palmieri *et al*., 2009; Moog *et al*., 2020).

## Funding Statement

SL acknowledges a China Scholarship Council doctoral studentship grant awarded via the Université Paris-Saclay, and SMY acknowledges a visiting student exchange grant, anchoring the Biology Centre CAS in the European Research Area (grant number: CZ.02.2.69/0.0/0.0/18_054/0014649) and GA JU travel award from the University of South Bohemia. CB acknowledges French Government ‘Investissements d’Avenir’ programs OCEANOMICS (ANR-11-BTBR-0008), FRANCE GENOMIQUE (ANR-10-INBS-09-08), MEMO LIFE (ANR-10-LABX-54), and PSL Research University (ANR-11-IDEX-0001-02), as well as funding from the European Research Council (ERC) under the European Union’s Horizon 2020 research and innovation program (Diatomic; grant agreement No. 835067), and the Agence Nationale de la Recherche (BrownCut project; ANR-19-CE20-0020). RGD acknowledges a CNRS Momentum Fellowship, awarded 2019-2021, and an ERC Starting Grant (“ChloroMosaic”, grant number 101039760, awarded 2023-2027).

## Author Contributions

RGD coordinated the study, performed phylogenetic and gene co-regulation analyses, synthesized gene constructs for biolistic transformation. SL performed biolistic transformations, visualized transformant lines of Phatr3_J42874 and Phatr3_J46612 with confocal microscopy, and performed all *Tara* Oceans analyses. SMY performed confocal microscopy visualization of Phatr3_J46742 and aided with Mitotracker staining. MO and CB acquired funding for the study, and provided essential and unpublished genetic resources for the gene co-regulation and *Tara* Oceans analyses performed. RGD wrote the manuscript with input from SMY and CB. All co-authors read, provided critical feedback and approved the manuscript prior to submission.

## Data availability statement

Raw confocal microscopy images, retrieved homologue sequences, and *Tara* Oceans distributions are provided in the linked osf.io repository (doi: 10.17605/OSF.IO/89VM3), in the subfolder “Transporters > MCF proteins”. All remaining supporting data, including alignments and phylogenies of cultured species sequences, are provided in **Tables S1-S4**. Pre-analyzed RNAseq data described in this study is available from the DiatomicBase server: https://www.diatomicsbase.bio.ens.psl.eu/. Cryopreserved plasmids and transformant lines used for confocal visualization of MCFc family proteins are available from the corresponding author on reasonable request.

---

**Fig. S1.**
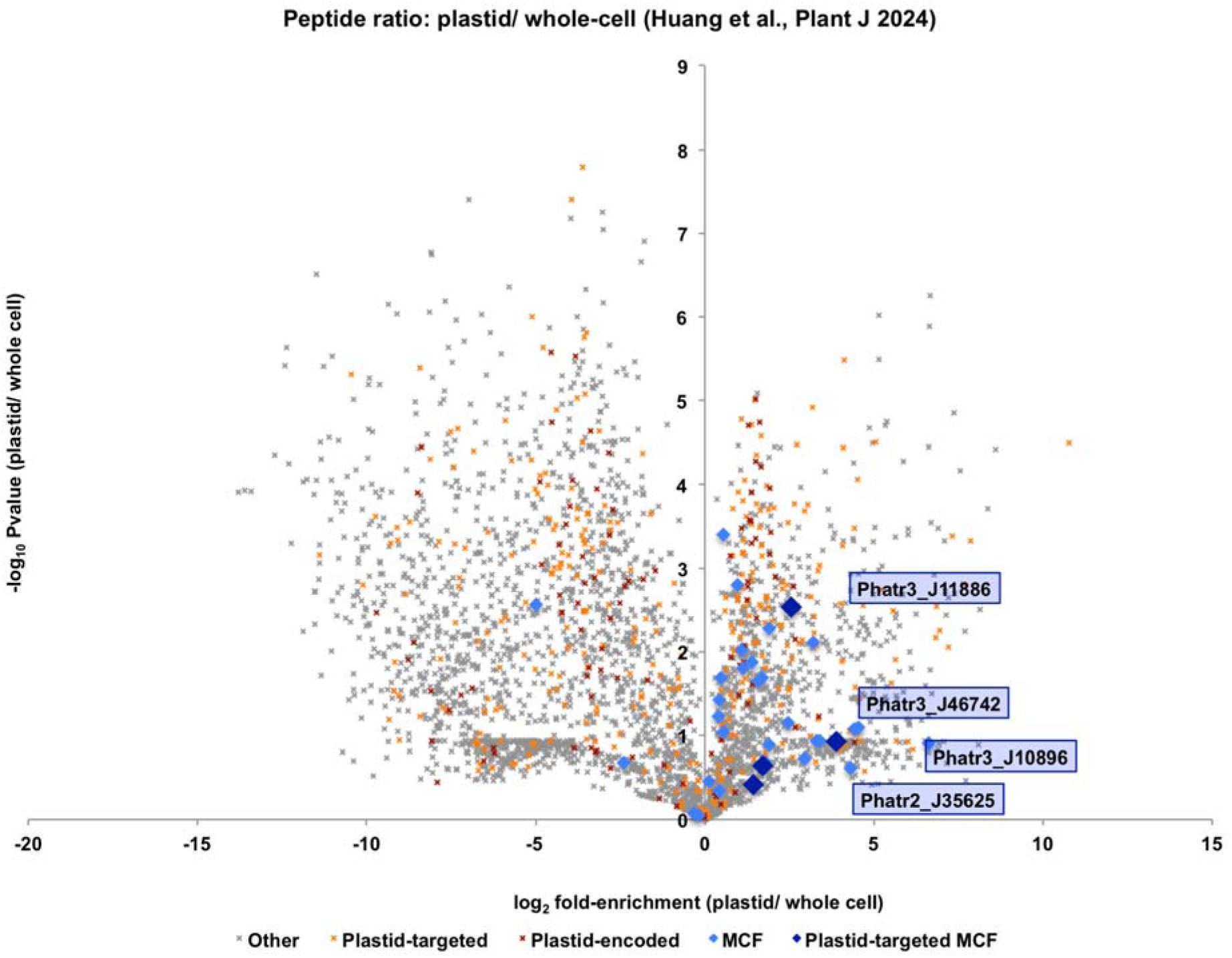
**Occurrence of MCF domains in plastid-enriched versus total cellular experimental proteomic fractions**, shown as per Huang et al. (2014). Proteins are shaded per encoded genome and predicted *in silica* localization, based on consensus ASAFind and HECTAR targeting prediction. Four MCF protein proteins with predicted plastid localization detectable both in plastid-enriched and total cell fractions are labelled, in each case showing enrichment in plastid fractions.

**Fig. S2.**
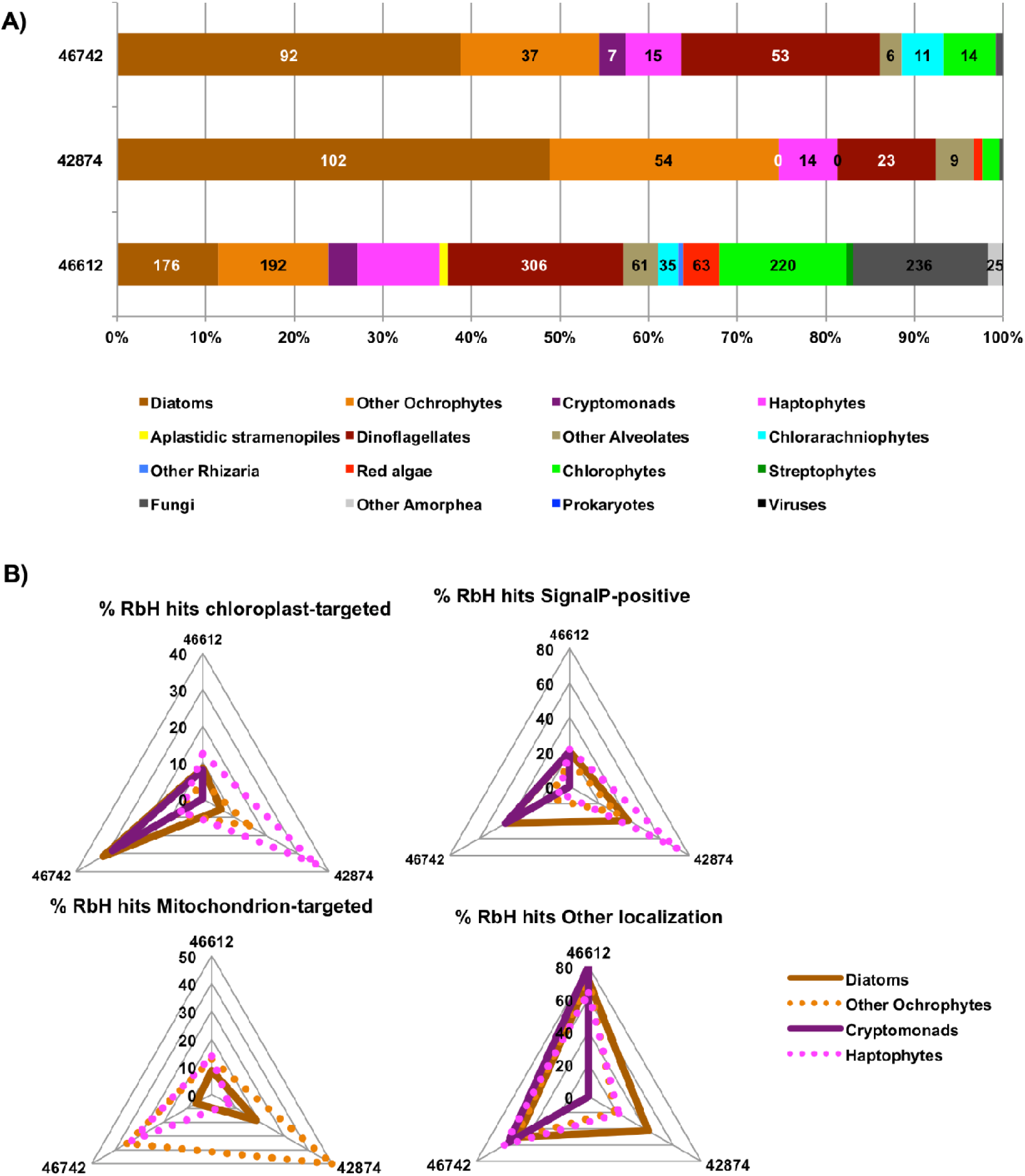
Distribution and localisation of homologues to Phatr3_J46742. **Panel A** shows the distribution of reciprocal BLAST best-hit matches (with threshold e-evalue 10’^05^) to MCFc/ Phatr3_J46742 and two related proteins (Phatr3_J42874, Phatr3_J466l2) encoded in the *Phaeodactylum tricornutum* genome (Villar et al., 2024). **Panel B** shows the proportion of these proteins for species with four-membrane bound chloroplasts (Åe., ochrophytes, cryptomonads and haptophytes) inferred to localise to the chloroplast (using ASAFiπd or HECTAR *in silico* prediction; Gschloessl etal., 2008; Gruber et al., 2015), predicted to contain an N-terminal signal peptide (using SignalP v 3.0; Bendtsen et al., 2005), predicted to encode a mitochondrial presequence (using HECTAR or MitoFates, Fukusawa et al. 2015) or with no predicted localization. Homologues of MCFc show a notable propensity towards chloroplast localizations in diatoms, whereas homologues of Phatr3_J42874 and Phatr3_J466l2 show more frequent endomembrane, mitochondrial or untargeted localization predictions.

**Fig. S3.**
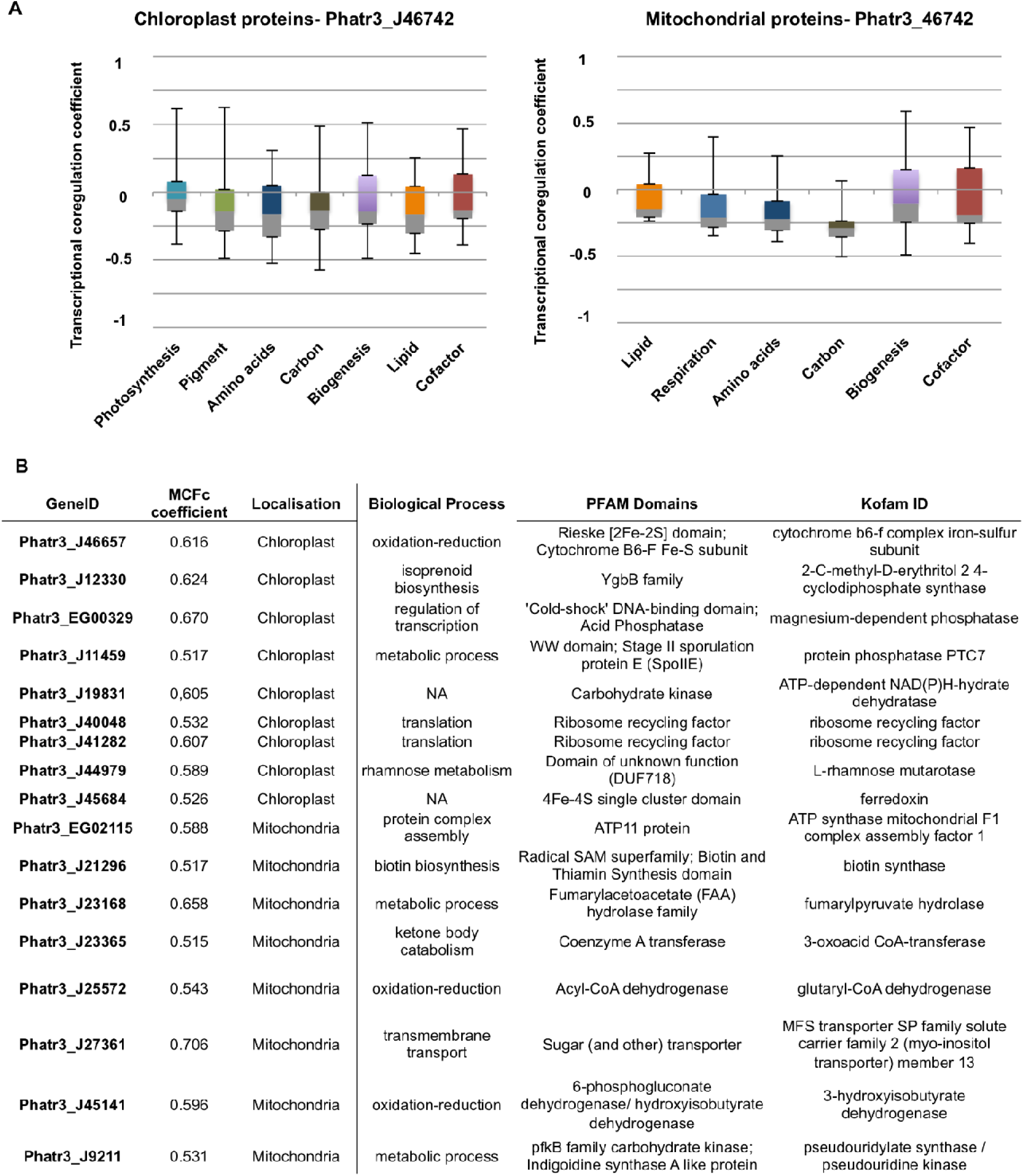
Identification of *P. trícornutum* genes coexpressed with Phatr3_J46742. **A:** boxplots of mean Spearman correlations of co-regulation of proteins associated with *P. tricornutum* organelle proteomes to Phatr3_J46742 in a meta-analysis of published microarray and transcriptome data (Ashworth et al., 2016; Villar et al., 2024) following the methodology of Liu et al. 2022. Phatr3_J46742 is typically found to be anticorrelated to most core organelle processes, with coregulation coefficients typically lower than −0.25 to most organelle-targeted proteins of annotated function. **B:** details of 17 *P. trícornutum* genes coding for organelle-targeted proteins with **(i)** functional annotations from combined PFAM and Kofam annotation, and **(ii)** co-regulation coefficients > 0.5 to MCFc. These notable encode a putative mitochondria-targeted major facilitator superfamily transporter (genelD: Phatr3_J 27 361) with co-regulation coefficient >0.7.

**Fig. S4.**
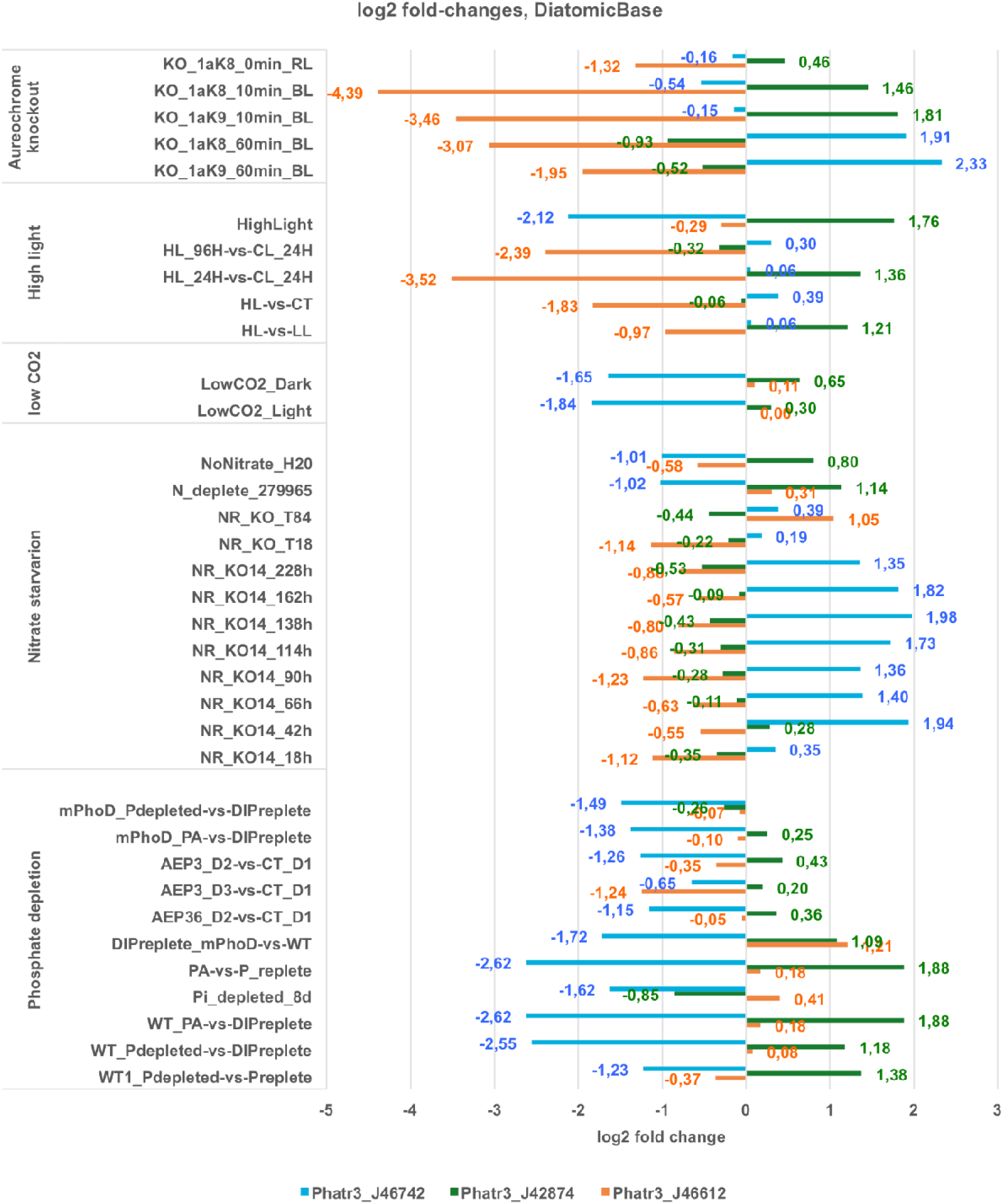
Transcriptional regulation of three related MCFc genes. This figure plots relative fold-changes deemed to be statistically significant (p < 0,05) in a meta-study of published *P. tricomutum* RNAseq data (Villar et al., 2024). Transcriptomic data are grouped by conditions. Briely, PhatrJ_466l2 is dowπregulated in aureochrome knockout, high light, and nitrate starvation conditions; Phatr3 J42784 is upregulated under short-term illumination in aureochrome knockouts, high light conditions, and phosphate depletion; and Phatr3_J46742 is upregulated under long-term illumination in aureochrome knockouts and nitrate starvation, but dowπregulated under low CO_2_ and phosphate depletion conditions.

**Fig. S5.**
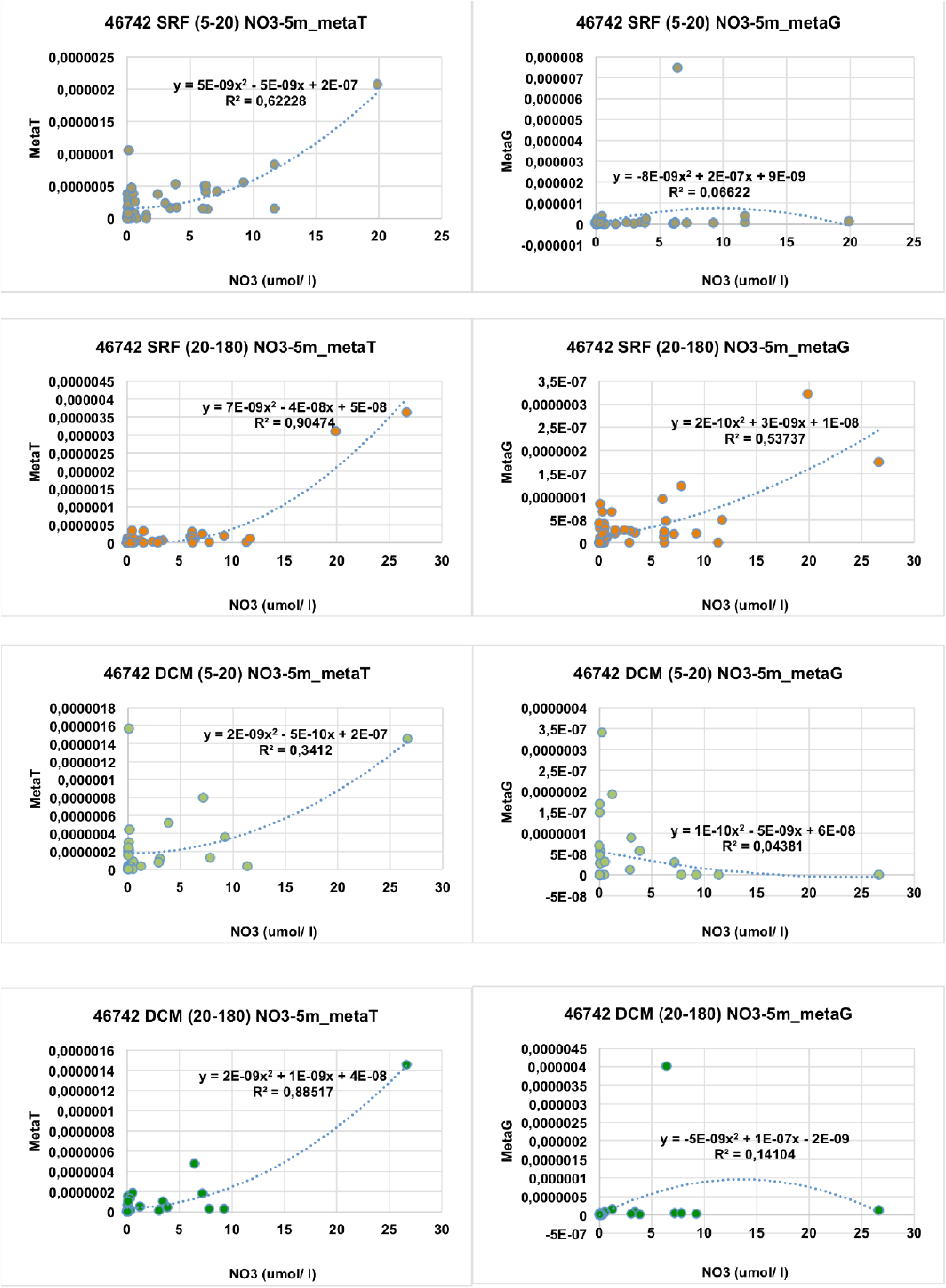
Scatterplots of Phatr3_J46742 metaT/G abundances and NO_3_^-^, 5m depth. Broadly, Phatr3_J46742 meta-traπscripts show strong positive correlations to nitrate abundance, where meta-geπes show no clear correlation.

**Fig. S6.**
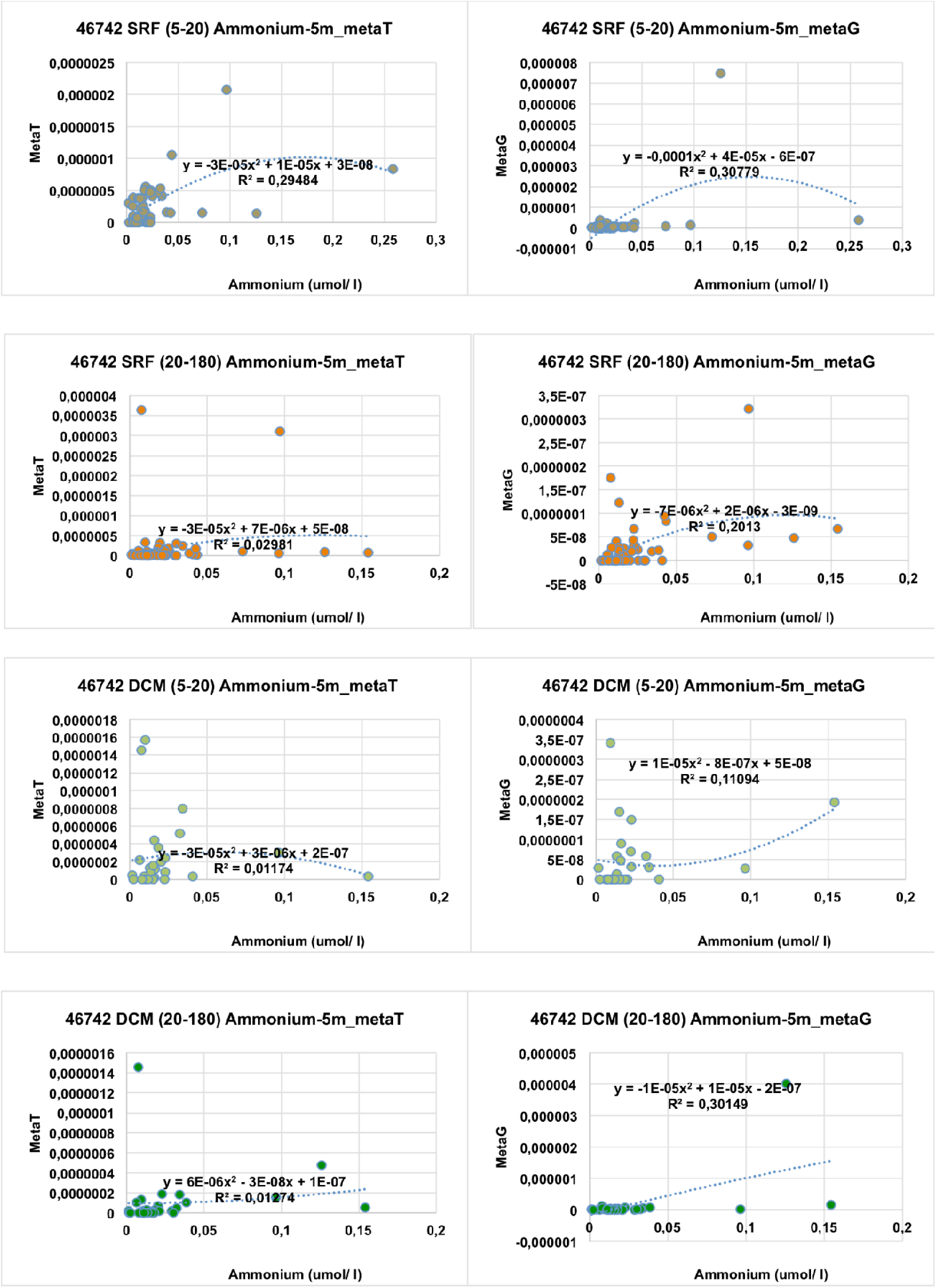
Scatterplots of Phatr3_J46742 metaT/G abundances and NH_4_^+^, 5m depth. No significant correlations are observed between ammonium and imetaT, despite (for DCM only) weak positive correlations to metaG abundance.

**Fig. S7.**
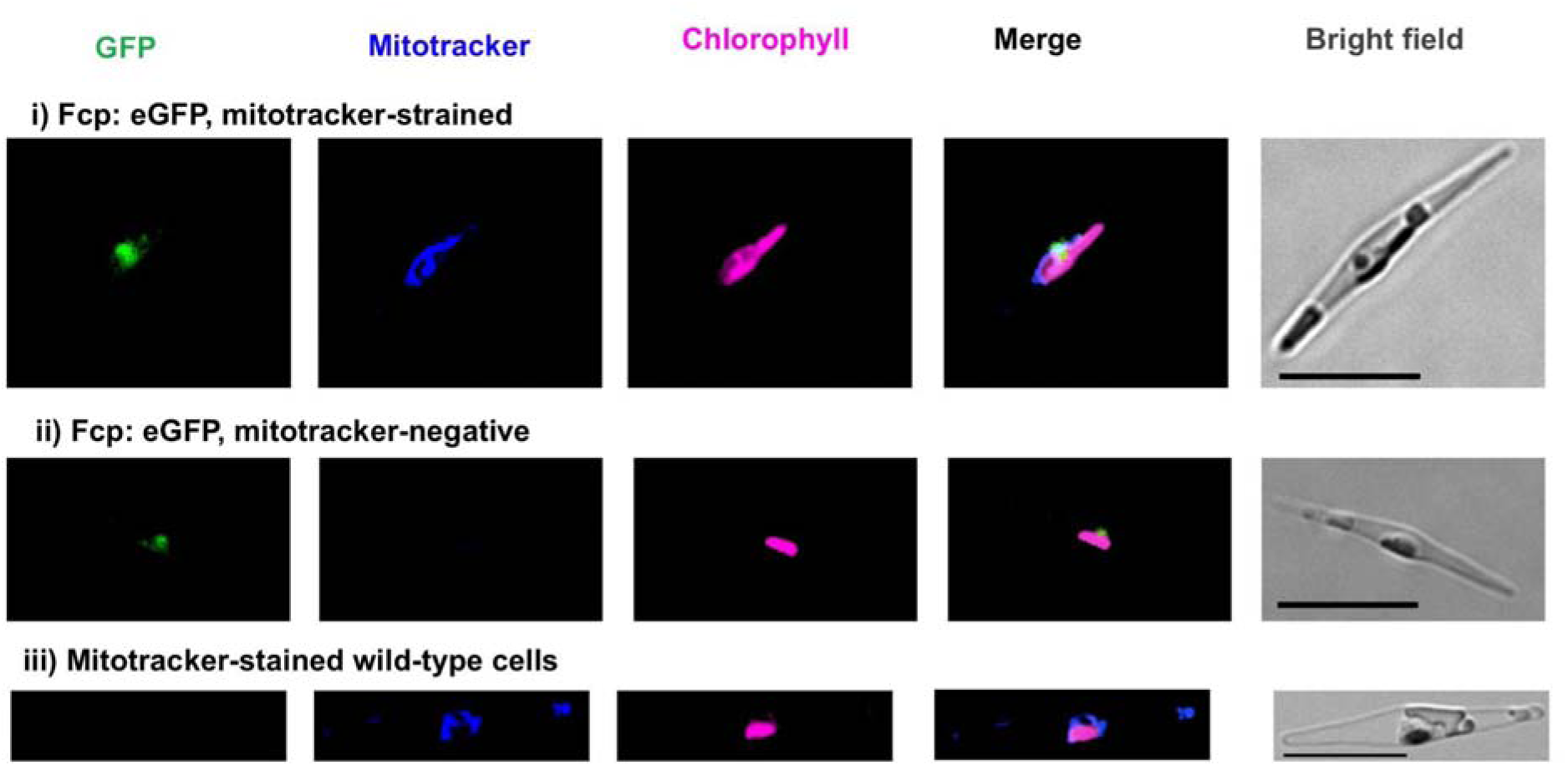
Control images for localization of *Phaeodactylum* MCFc homologues. This figure shows confocal microscopy images of transgenic *P. tricornutum* expressing C-terminal linked GFP constructs stained and unstained cell lines expressing unlinked cytoplasmic GFP, and Mitotracker-stained wild-type cell lines, as per Fig. 3.

## Acknowledgments

This paper is contribution *XXX* to Tara Oceans. The authors acknowledge Priscillia Pierre-Elies (Institut de Biologie de l’Ecole Normale Supérieure) and Pauline Clémente (Lycée ENCPB-Gilles de Gennes) for aid with the biolistics transformation and the confocal microscopy shown in **Fig. 3**.

## Notes

https://osf.io/89vm3/

### Competing Interest Statement

The authors have declared no competing interest.

