## Supplementary material for "Dynamic relocalization and divergent expression of a major facilitator carrier subfamily in diatoms": Figures S1-7

### Supporting Figures and Supporting Table Legends

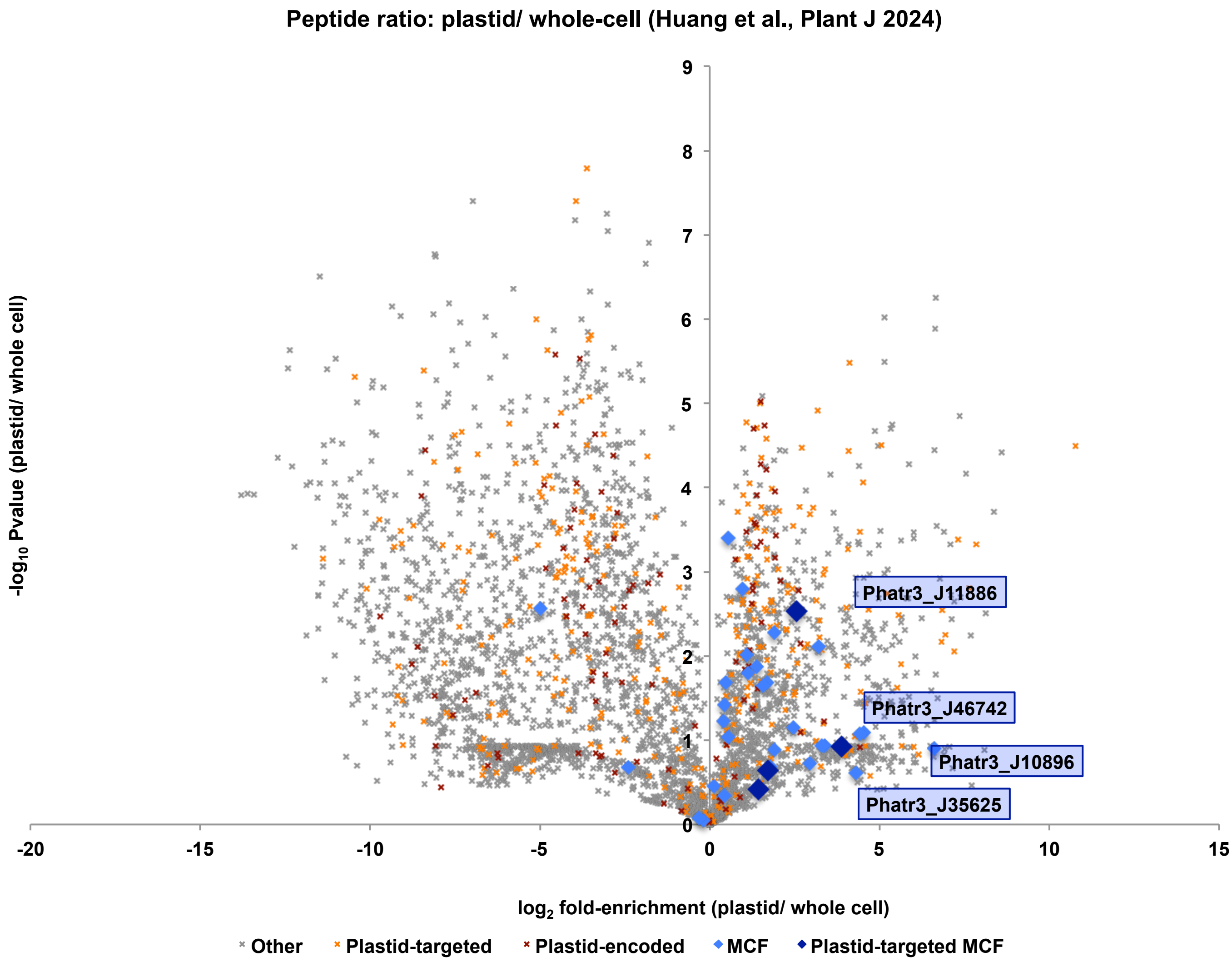


**Fig. S1. Occurrence of MCF domains in plastid-enriched versus total cellular experimental proteomic fractions,** shown as per Huang et al. (2014). Proteins are shaded per encoded genome and predicted *in silico* localization, based on consensus ASAFind and HECTAR targeting prediction. Four MCF protein proteins with predicted plastid localization detectable both in plastid-enriched and total cell fractions are labelled, in each case showing enrichment in plastid fractions.

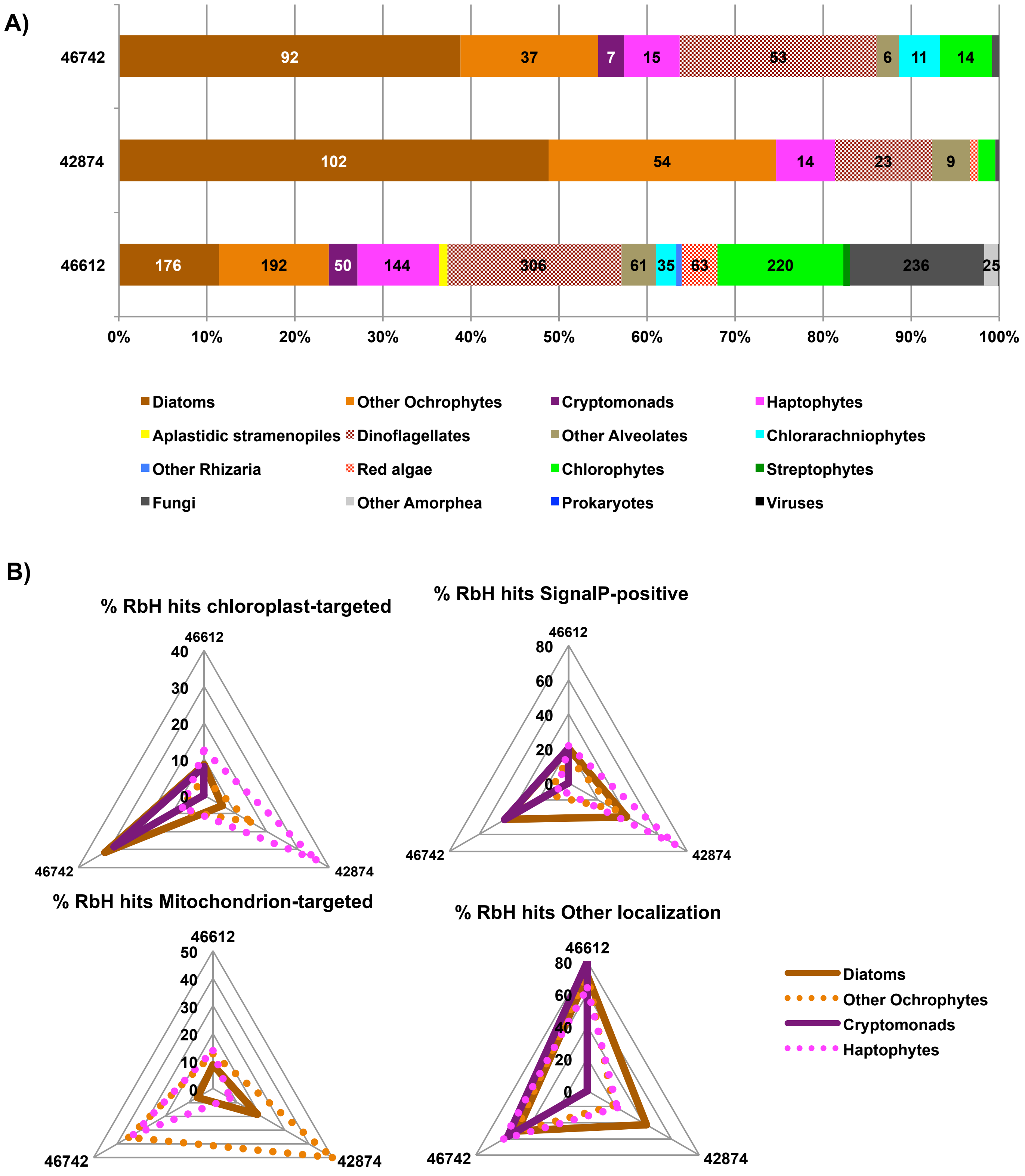


**Fig. S2. Distribution and localisation of homologues to Phatr3_J46742. Panel A** shows the distribution of reciprocal BLAST best-hit matches (with threshold e-evalue $10^{-05}$) to MCFc/ Phatr3_J46742 and two related proteins (Phatr3_J42874, Phatr3_J46612) encoded in the *Phaeodactylum tricornutum* genome (Villar et al., 2024). **Panel B** shows the proportion of these proteins for species with four-membrane bound chloroplasts (*i.e.,* ochrophytes, cryptomonads and haptophytes) inferred to localise to the chloroplast (using ASAFind or HECTAR *in silico* prediction; Gschloessl et al., 2008; Gruber et al., 2015), predicted to contain an N-terminal signal peptide (using SignalP v 3.0; Bendtsen et al., 2005), predicted to encode a mitochondrial presequence (using HECTAR or MitoFates, Fukusawa et al. 2015) or with no predicted localization. Homologues of MCFc show a notable propensity towards chloroplast localizations in diatoms, whereas homologues of Phatr3_J42874 and Phatr3_J46612 show more frequent endomembrane, mitochondrial or untargeted localization predictions.

**A**

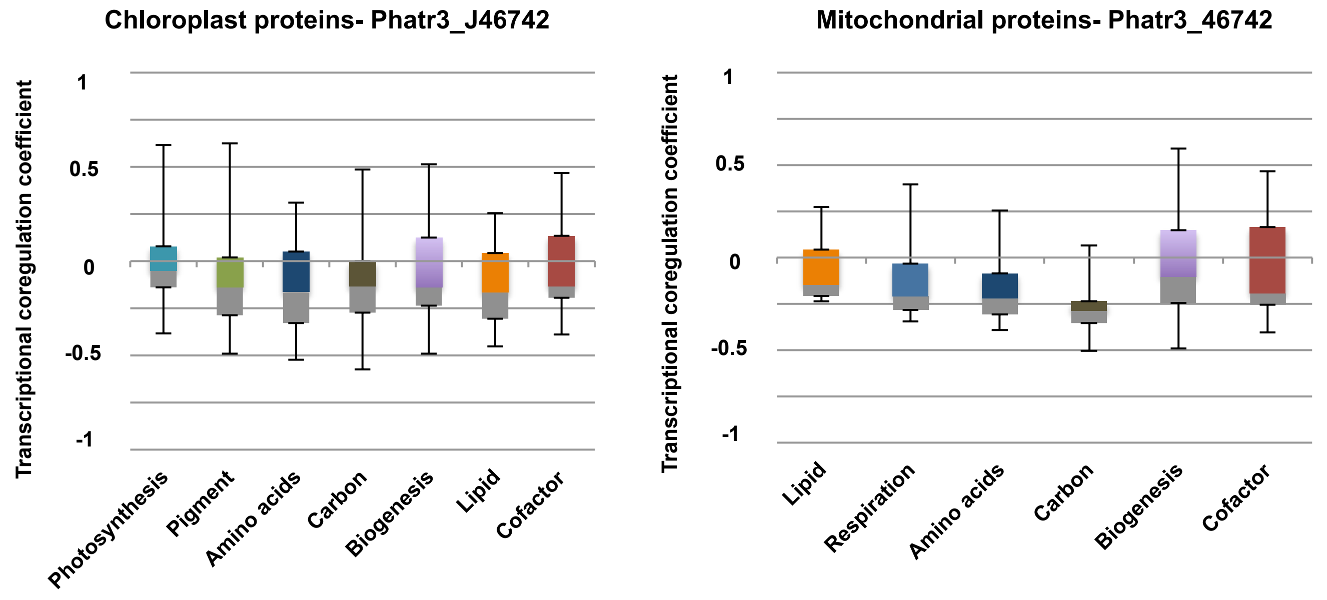


**B**

| GeneID | MCFc coefficient | Localisation | Biological Process | PFAM Domains | Kofam ID |
|---|---|---|---|---|---|
| **Phatr3_J46657** | 0.616 | Chloroplast | oxidation-reduction | Rieske [2Fe-2S] domain; Cytochrome B6-F Fe-S subunit | cytochrome b6-f complex iron-sulfur subunit |
| **Phatr3_J12330** | 0.624 | Chloroplast | isoprenoid biosynthesis | YgbB family | 2-C-methyl-D-erythritol 2 4-cyclodiphosphate synthase |
| **Phatr3_EG00329** | 0.670 | Chloroplast | regulation of transcription | 'Cold-shock' DNA-binding domain; Acid Phosphatase | magnesium-dependent phosphatase |
| **Phatr3_J11459** | 0.517 | Chloroplast | metabolic process | WW domain; Stage II sporulation protein E (SpoIIE) | protein phosphatase PTC7 |
| **Phatr3_J19831** | 0,605 | Chloroplast | NA | Carbohydrate kinase | ATP-dependent NAD(P)H-hydrate dehydratase |
| **Phatr3_J40048** | 0.532 | Chloroplast | translation | Ribosome recycling factor | ribosome recycling factor |
| **Phatr3_J41282** | 0.607 | Chloroplast | translation | Ribosome recycling factor | ribosome recycling factor |
| **Phatr3_J44979** | 0.589 | Chloroplast | rhamnose metabolism | Domain of unknown function (DUF718) | L-rhamnose mutarotase |
| **Phatr3_J45684** | 0.526 | Chloroplast | NA | 4Fe-4S single cluster domain | ferredoxin |
| **Phatr3_EG02115** | 0.588 | Mitochondria | protein complex assembly | ATP11 protein | ATP synthase mitochondrial F1 complex assembly factor 1 |
| **Phatr3_J21296** | 0.517 | Mitochondria | biotin biosynthesis | Radical SAM superfamily; Biotin and Thiamin Synthesis domain | biotin synthase |
| **Phatr3_J23168** | 0.658 | Mitochondria | metabolic process | Fumarylacetoacetate (FAA) hydrolase family | fumarylpyruvate hydrolase |
| **Phatr3_J23365** | 0.515 | Mitochondria | ketone body catabolism | Coenzyme A transferase | 3-oxoacid CoA-transferase |
| **Phatr3_J25572** | 0.543 | Mitochondria | oxidation-reduction | Acyl-CoA dehydrogenase | glutaryl-CoA dehydrogenase |
| **Phatr3_J27361** | 0.706 | Mitochondria | transmembrane transport | Sugar (and other) transporter | MFS transporter SP family solute carrier family 2 (myo-inositol transporter) member 13 |
| **Phatr3_J45141** | 0.596 | Mitochondria | oxidation-reduction | 6-phosphogluconate dehydrogenase/ hydroxyisobutyrate dehydrogenase | 3-hydroxyisobutyrate dehydrogenase |
| **Phatr3_J9211** | 0.531 | Mitochondria | metabolic process | pfkB family carbohydrate kinase; Indigoidine synthase A like protein | pseudouridylate synthase / pseudouridine kinase |

**Fig. S3. Identification of *P. tricornutum* genes coexpressed with Phatr3_J46742 A:** boxplots of mean Spearman correlations of co-regulation of proteins associated with *P. tricornutum* organelle proteomes to Phatr3_J46742 in a meta-analysis of published microarray and transcriptome data (Ashworth et al., 2016; Villar et al., 2024) following the methodology of Liu et al. 2022. Phatr3_J46742 is typically found to be anticorrelated to most core organelle processes, with coregulation coefficients typically lower than -0.25 to most organelle-targeted proteins of annotated function. **B:** details of 17 *P. tricornutum* genes coding for organelle-targeted proteins with **(i)** functional annotations from combined PFAM and Kofam annotation, and **(ii)** co-regulation coefficients > 0.5 to MCFc. These notable encode a putative mitochondria-targeted major facilitator superfamily transporter (geneID: Phatr3_J27361) with co-regulation coefficient >0.7.

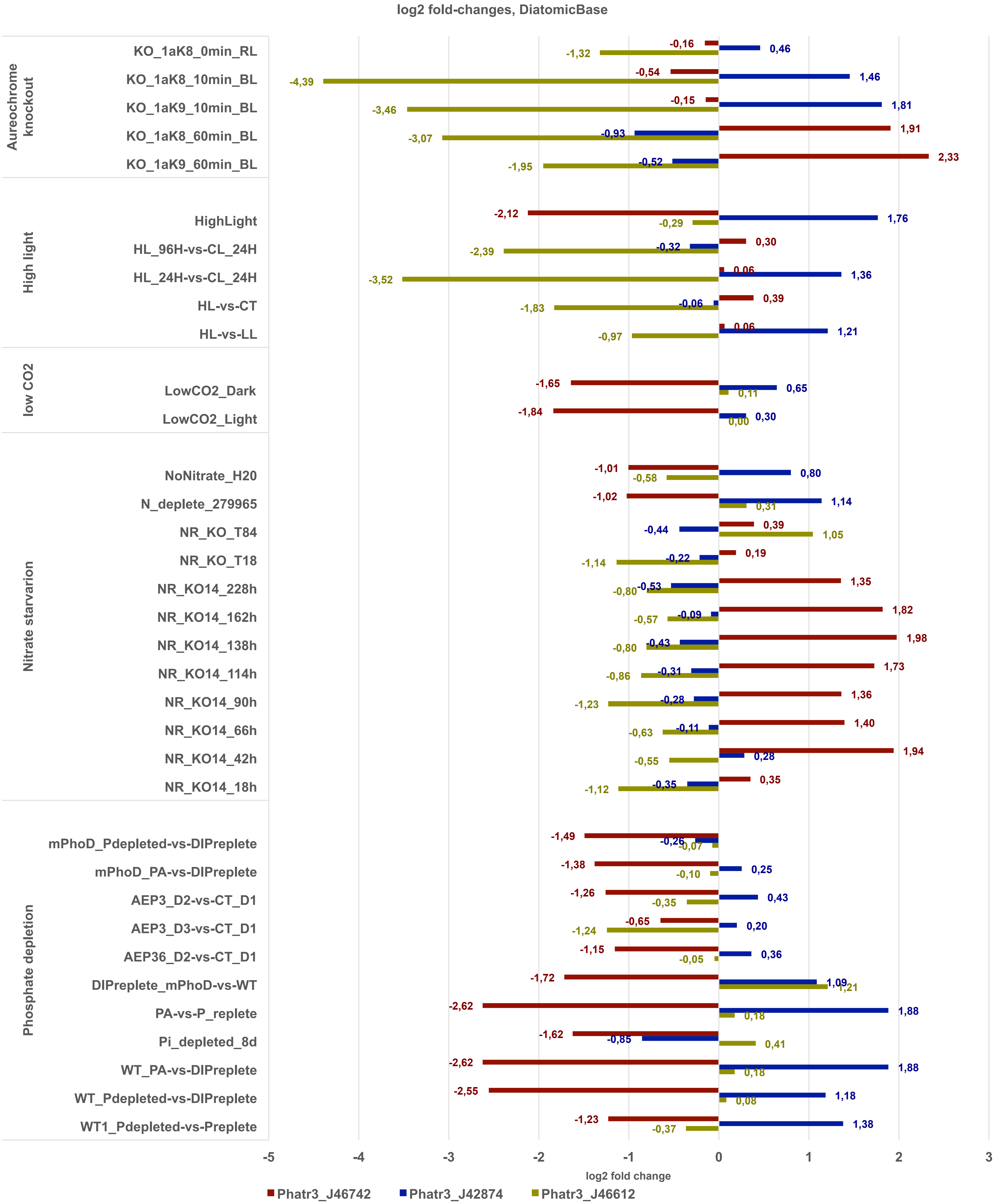


**Fig. S4. Transcriptional regulation of three related MCFc genes.** This figure plots relative fold-changes deemed to be statistically significant ($p < 0.05$) in a meta-study of published *P. tricornutum* RNAseq data (Villar et al., 2024). Transcriptomic data are grouped by conditions. Briely, PhatrJ_46612 is downregulated in aureochrome knockout, high light, and nitrate starvation conditions; Phatr3_J42784 is upregulated under short-term illumination in aureochrome knockouts, high light conditions, and phosphate depletion; and Phatr3_J46742 is upregulated under long-term illumination in aureochrome knockouts and nitrate starvation, but downregulated under low $CO_2$ and phosphate depletion conditions.

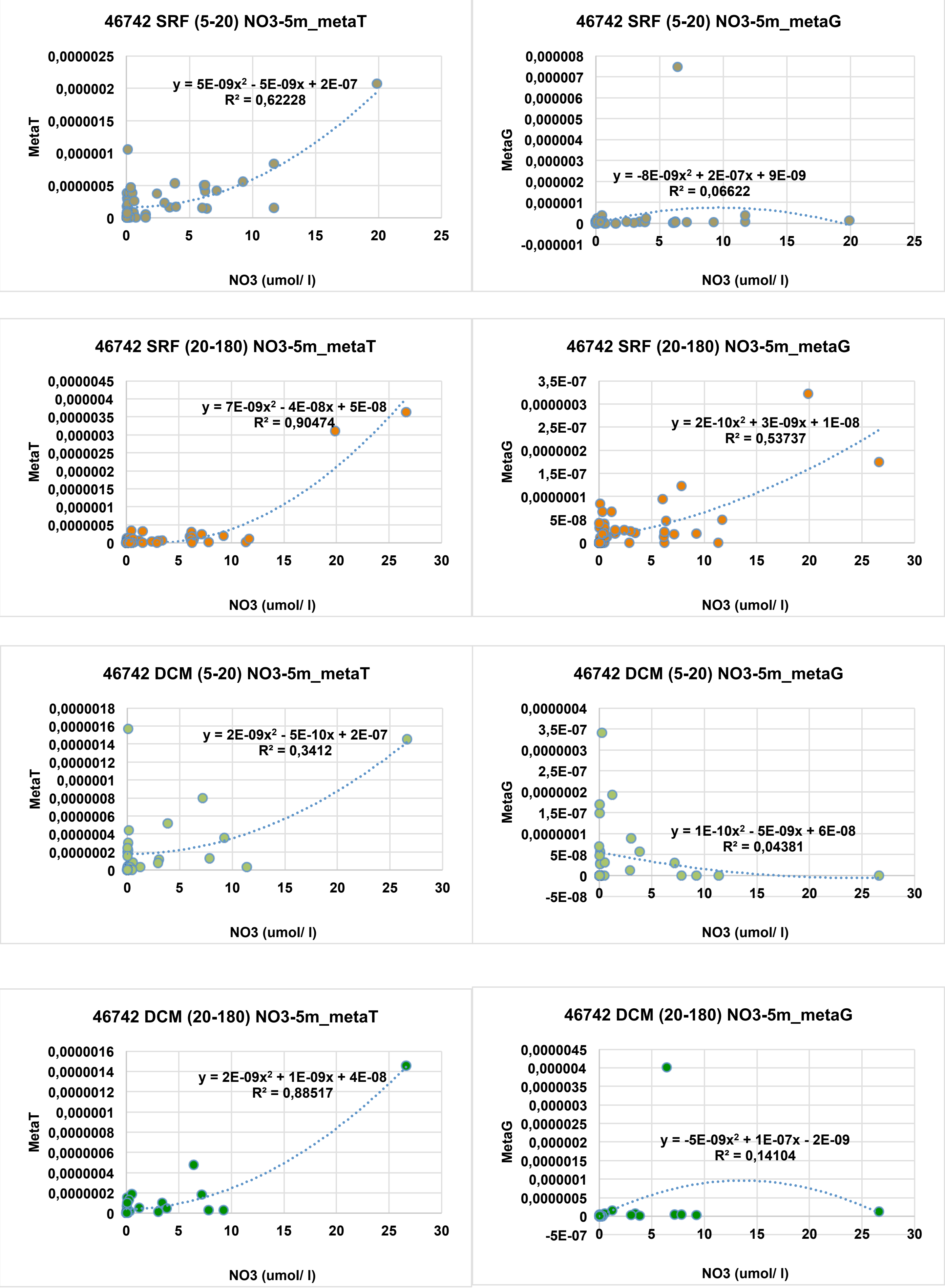


**Fig. S5. Scatterplots of Phatr3_J46742 metaT/G abundances and simulated $NO_3^-$, 5m depth.** Phatr3_J46742 meta-transcripts show strong positive correlations to nitrate abundance, where meta-genes show no clear correlation.

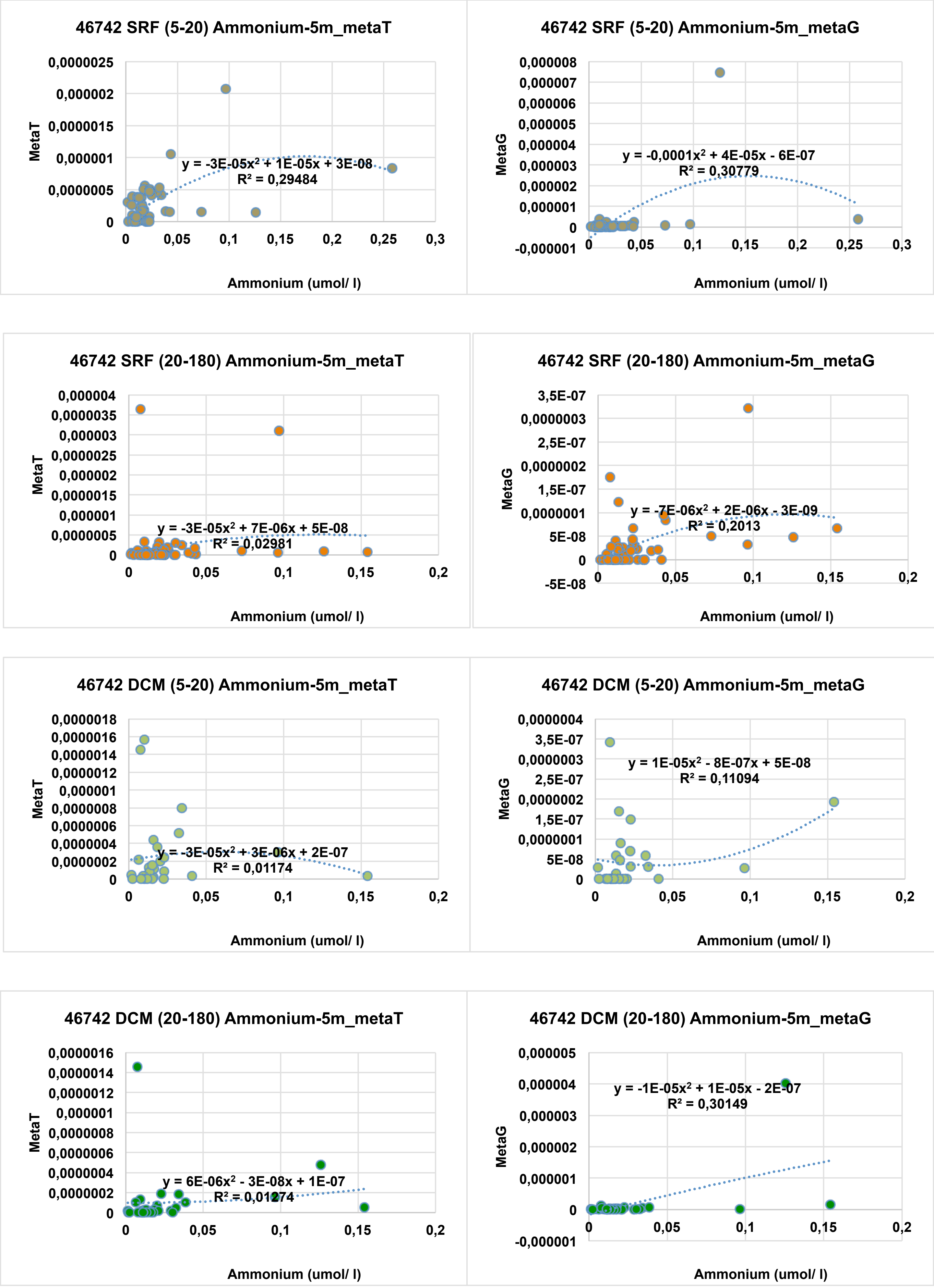


**Fig. S6. Scatterplots of Phatr3_J46742 metaT/G abundances and simulated $NH_4^+$, 5m depth.** No significant correlations are observed between ammonium and metaT, despite weak positive correlations to metaG abundance.

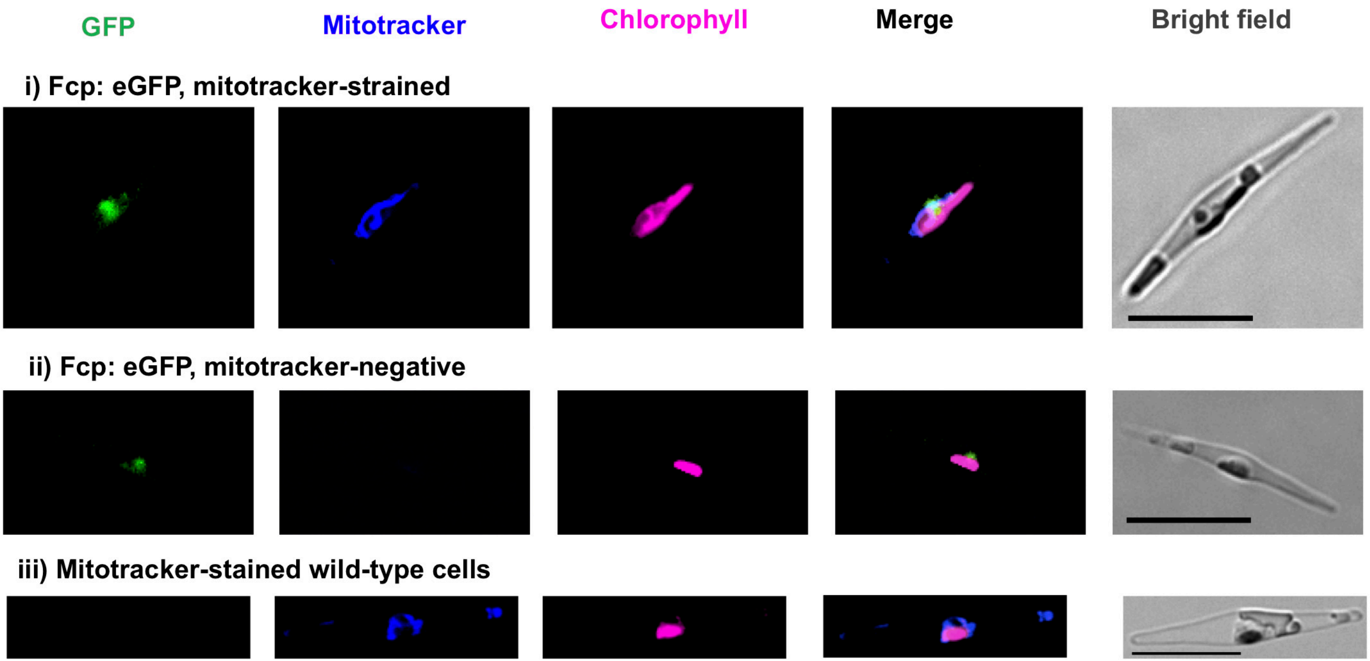


**Fig. S7. Control images for localization of *Phaeodactylum* MCFc homologues.** This figure shows confocal microscopy images of transgenic *P. tricornutum* expressing C-terminal linked GFP constructs stained and unstained cell lines expressing unlinked cytoplasmic GFP, and Mitotracker-stained wild-type cell lines, as per **Fig. 3**.

**Table S1. Phylogeny of MCFc and its homologues across the algal tree of life.**

This workbook provides sequences, alignments, and phylogenetic topologies of proteins identified to be homologous to MCFc or to two other closely related MCF transporters (Phatr3_J42874, Phatr3_J46612) encoded in the *Phaeodactylum tricornutum* version 3 genome annotation.

The first sheet ("**MCF matrix**") tabulates all MCF domains in the Phaeodactylum genome, their predicted KEGG annotations and localizations, and the e-values of their reciprocal BLASTp similarity to one another.

The second sheet ("**Ingroups**") lists reciprocal BLASTp best-hit homologues to each protein, identified with threshold e-value 10-05, from 289 different diatom, ochrophyte (photosynthetic diatom relative), non-photosynthetic stramenopile (diatom relative), cryptomonad and haptophyte (other groups bearing four-membraned plastid) genomes and transcriptomes, following data from Liu et al, 2022. The ID and sequence of each homologue are listed, alongside their best-scoring reciprocal BLASTp hit against the complete version 3 *Phaeodactylum* genome.

The third sheet ("**Outgroups**") lists equivalent homologues identified from a combined reference library of uniprot, jgi genomes, and MMETSP transcriptomes from across the tree of life, and validated by reciprocal BLASTp best hit analysis.

The fourth sheet ("**Trimmed alignment**") provides a nexus format alignment of a subselected 78 taxa x 262 aa set of homologues, prioritising homologues with clear N-terminal targeting sequences to the plastids, mitochondria or other cytoplasmic organelles, taxonomic balance, and also sites with sufficient (<20% gapped) conservation to be phylogenetically informative.

The fifth sheet ("**Trees**") provides consensus MrBayes topologies realised with three substitution matrices (GTR, Jones, WAG) and RAxML with three matrices (GTR, JTT, WAG) for the trimmed alignment.

The final sheet ("**RbH heatmap**") tabulates the total number of homologues from different taxonomic groups identified by reciprocal BLASTp best-hit analysis, alongside their consensus localizations (chloroplast, endomembrane-targeted, mitochondrion-targeted, other) for lineages with four-membraned plastids (diatoms, other ochrophytes, haptophytes, cryptomonads).

**Table S2. Primers and constructs used for GFP localization of MCF (Phatr3_J46742) and two related proteins (Phatr3_J46612, Phatr3_J42874).**

This table provides **(top)** primer sequences and coding constructs and **(bottom)** an exemplar plasmid map for pPhat Fcp: C-terminal eGFP localization constructs generated. Plasmids were synthesized using Gibson cloning with primers fused (Gibson 5', Gibson 3') either to the start and end of the coding sequence minus the stop codon (Phatr3_J42874, Phatr3_J46612) or to the final 209 bp of the 5' UTR and the first 87 aa of the encoded protein (Phatr3_J46742), avoiding out-of-frame termination codons. Constructs were fused to a linear pPhat-eGFP vector amplified with complementary primers to the FcpA promoter and to the eGFP region minus the initiator methionine.

**Table SW. Identification of genes co-expressed with MCFc in the *P. tricornutum***

**genome.**

This table provides co-regulation coefficients (Spearman correlations of gene expression relative abundance) for MCFc/ Phatr3_J46742 and genes coding for two closely related MCF transporters (Phatr3_J42874, Phatr3_J46612) in a meta-analysis of previously published microarray and RNAseq data for *P. tricornutum*, following the methodology of Liu et al., 2022.

Sheet 1 ("**Phat3 genome**") provides a tabulated gene list for P. tricornutum alongside consensus protein localization, epigenetic and functional (PFAM, KEGG) annotations per Aït-Mohamed et al., 2020. Columns E**-G** show the co-regulation coefficients calculated for all three genes coding for MCF transporters related to MCFc.

Sheets 2 and 3 ("**Organelle metabolism**", "**Non-redundant organelle metabolism**") tabulate co-regulation coefficients for annotated core chloroplast and mitochondrial metabolic pathways against each MCF gene, following methodology of Aït-Mohamed et al., 2020.

Sheets 4, 6 and 8 ("**46472/ 42874/ 46612 organelle ANOVAs**") present boxplots and one-way analyses of variance of the difference in the mean co-regulation coefficients calculated for genes encoding specific functional groups of chloroplast- or mitochondria-targeted proteins, compared to those of all other genes encoding proteins with the same localization. P-values are signed such that positive values indicate the mean is greater than expected, and negative values that it is lower than expected.

Sheets 5, 7 and 9 ("**46472/ 42874/ 46612 organelle map**") presents schematic maps of core Phaeodactylum organelle proteomes, adapted from Ait-Mohamed et al., 2020, and shaded by transcriptional coregulation of the underlying gene to genes encoding MCF proteins. A scale bar is provided to the right of each map.

**Table S4. Relative abundances of diatom MCF family transporter meta-genes in *Tara* Oceans data.**

This workbook provides data regarding the relative mapped abundances (expressed per 1 = all reads for each sample) for meta-genes corresponding to three closely-related diatom MCF transporters (Phatr3_J46742/ MCFc- chloroplastic, Phatr3_J46612- mitochondrial, Phatr3_J42874- endomembrane) identified from *Tara* meta-genes.

Sheets 1-6 (**"46742-metaT", "46742-metaG", "46612-metaT", "46612-metaG", "42874-metaT", "42874-metaG"**) provide total mapped relative abundances for all meta-genes phylogenetically reconciled to be of diatom origin and to resolve to each MCF sub-family, following the methodology of Liu et al., 2022.

Sheet 7 ("**Totals**") provide the total relative mapped abundances for each MFC sub-family for each depth and size-fraction. Values are provided both for absolute and ranked values within each depth/ size-fraction combinations, alongside sum-differences between the metaT and metaG relative abundances, considering both absolute total relative abundances and rank values.

Sheets 8-10 ("**Absolute correlations**", "**Ranked correlations**", "**Sum correlations**") provide correlation coefficients between the relative abundance of each MCF subfamily and each environmental variable measured over *Tara* stations, either considering the absolute

(Pearson) and ranked (Spearman) correlations of metaT and metaG separately, or the sum (difference) between metaT and metaG values. Data are provided for 5-20 um and 20-180 um size fractions, and SRF and DCM depths.

Sheets 11-12 ("**Correlation tests**", "**Correlation heatmap**") provide raw and synthesized statistics for parametric test comparisons of the correlation of MCF family meta-transcript and meta-gene abundances to each environmental variable, given the dependent relationships between metaT and metaG data observed across the entire dataset. Data are provided for 5-20 and 20-180 um size fractions, and surface and DCM depths.

Sheet 13 ("**Scatterplots**") show the different relationships of MCFc/ Phatr3_J46742 meta-gene total relative abundance in metaT and metaG data to measured nitrate and ammonium concentrations at 5m depth, for 5-20 and 20-180 um size fractions over surface and DCM depths.
